# Divergent mesolimbic dopamine circuits support alcohol-seeking triggered by discrete cues and contexts

**DOI:** 10.1101/475343

**Authors:** M.D. Valyear, I. Glovaci, A. Zaari, S. Lahlou, I. Trujillo-Pisanty, C.A. Chapman, N. Chaudhri

**Author notes:** Corresponding Author: Nadia Chaudhri, PhD, CSBN/GRNC, Department of Psychology, Concordia University, 7141 Sherbrooke Street West, Room SP 244, Montreal, QC, H4B-1R6, Canada.

## Abstract

Discrete and contextual cues that predict alcohol trigger alcohol-seeking. However, the extent to which context influences alcohol-seeking triggered by discrete cues, and the neural mechanisms underlying these responses, are not well known. We show that, relative to a neutral context, a context associated with alcohol persistently elevated alcohol-seeking triggered by a discrete cue, and supported higher levels of priming-induced reinstatement. Alcohol-seeking triggered by a discrete cue in a neutral context was reduced by designer receptor-mediated inhibition of ventral tegmental area (VTA) dopamine neurons in TH::Cre rats. Inhibiting terminals of VTA dopamine neurons in the nucleus accumbens (NAc) core reduced alcohol-seeking triggered by a discrete cue, irrespective of context, whereas inhibiting VTA dopamine terminals in the NAc shell selectively reduced the elevation of alcohol-seeking triggered by a discrete cue in an alcohol context. This dissociation highlights unique roles for divergent mesolimbic dopamine circuits in alcohol-seeking driven by discrete and contextual environmental cues.

---

Environmental stimuli that accompany drug use exert considerable control over drug-seeking and drug-taking^1,2,3^. This control is the product of Pavlovian conditioning, through which discrete and contextual stimuli become established as cues that predict the unconditioned properties of drugs. Discrete cues associated with alcohol are temporally delimited signals that occur in close temporal proximity with alcohol consumption (e.g., the taste or smell of alcohol)^4,5^. Current knowledge about processes that underlie the impact of environmental stimuli on alcohol-seeking is principally derived from research that has focused on discrete cues^6,7^. In contrast, contexts associated with alcohol are stable configurations of multi-modal stimuli that occur in the background during alcohol use (e.g., the ambience of a bar)^8^. Alcohol contexts can trigger alcohol-craving^9^ and motivate the pursuit of alcohol^2^. Context can also control nearly every feature – the magnitude, timing and valence – of a conditioned response triggered by a discrete cue^10^. Thus, the interplay between discrete cues and alcohol contexts may be a crucial determinant of alcohol-seeking and relapse.

The nucleus accumbens (NAc) core and shell subregions of the ventral striatum are hypothesized to be differentially involved in drug-seeking triggered by discrete cues and contexts^11,12^. However, few studies have delineated the precise contributions of discrete cues and contexts to drug-seeking behaviour or the molecular and anatomical identity of the circuits that underpin these behaviours. Using an animal model, we show here that contemporaneous exposure to a discrete cue for alcohol in an alcohol-associated context produces an enduring elevation in alcohol-seeking, as well as heightened alcohol-seeking in a relapse test. Using pharmacological and chemogenetic approaches, we demonstrate the necessity of dopamine neurotransmission and of ventral tegmental area (VTA) dopamine neurons in alcohol-seeking triggered by a discrete cue, in isolation from contextual alcohol cues. Critically, we identify a point of separability in VTA dopaminergic circuits that support alcohol-seeking triggered by a discrete cue and the elevation of this behavior in an alcohol context. We show that VTA dopamine inputs to the NAc core are critical for alcohol-seeking triggered by a discrete cue irrespective of the context in which that cue is experienced, whereas dopamine inputs to the NAc shell are engaged to elevate alcohol-seeking triggered by a discrete cue in an alcohol context. Lastly, we confirm that chemogenetic actions on dopamine terminals modulate the activity of post-synaptic medium spiny neurons in the NAc. These data are the first to identify the molecular and anatomical nature of neural circuits that underlie the separable involvement of the NAc core and shell subregions in alcohol-seeking controlled by discrete cues and contexts.

## Results

### An alcohol context has an enduring impact on alcohol-seeking triggered by a discrete cue and amplifies relapse

We previously found that alcohol-seeking triggered by a discrete conditioned stimulus (CS) was significantly elevated in a context associated with prior alcohol intake (alcohol context), relative to a familiar, neutral context in which alcohol was never consumed^4,5^. To extend the clinical relevance of this finding, here we tested the longevity of this context effect, as well as the impact of an alcohol context on CS-triggered alcohol-seeking in a relapse model.

Outbred, male Long-Evans rats (n=22) were acclimated to drinking 15% ethanol in their home-cages (**Fig. S1a**) and then given Pavlovian conditioning sessions in which presentations of a discrete, auditory CS were paired with 15% ethanol (EtOH) that was delivered into a fluid port for oral intake. These sessions occurred in a distinct ‘alcohol context’ created by visual, olfactory and tactile stimuli in the conditioning chamber, and were alternated daily with sessions in a different, ‘neutral context’ where alcohol was never available (**Fig. 1a**). Rats readily learned to selectively check the fluid port for alcohol during the CS (**Fig. S2a and S3a**).

**Fig 1.**
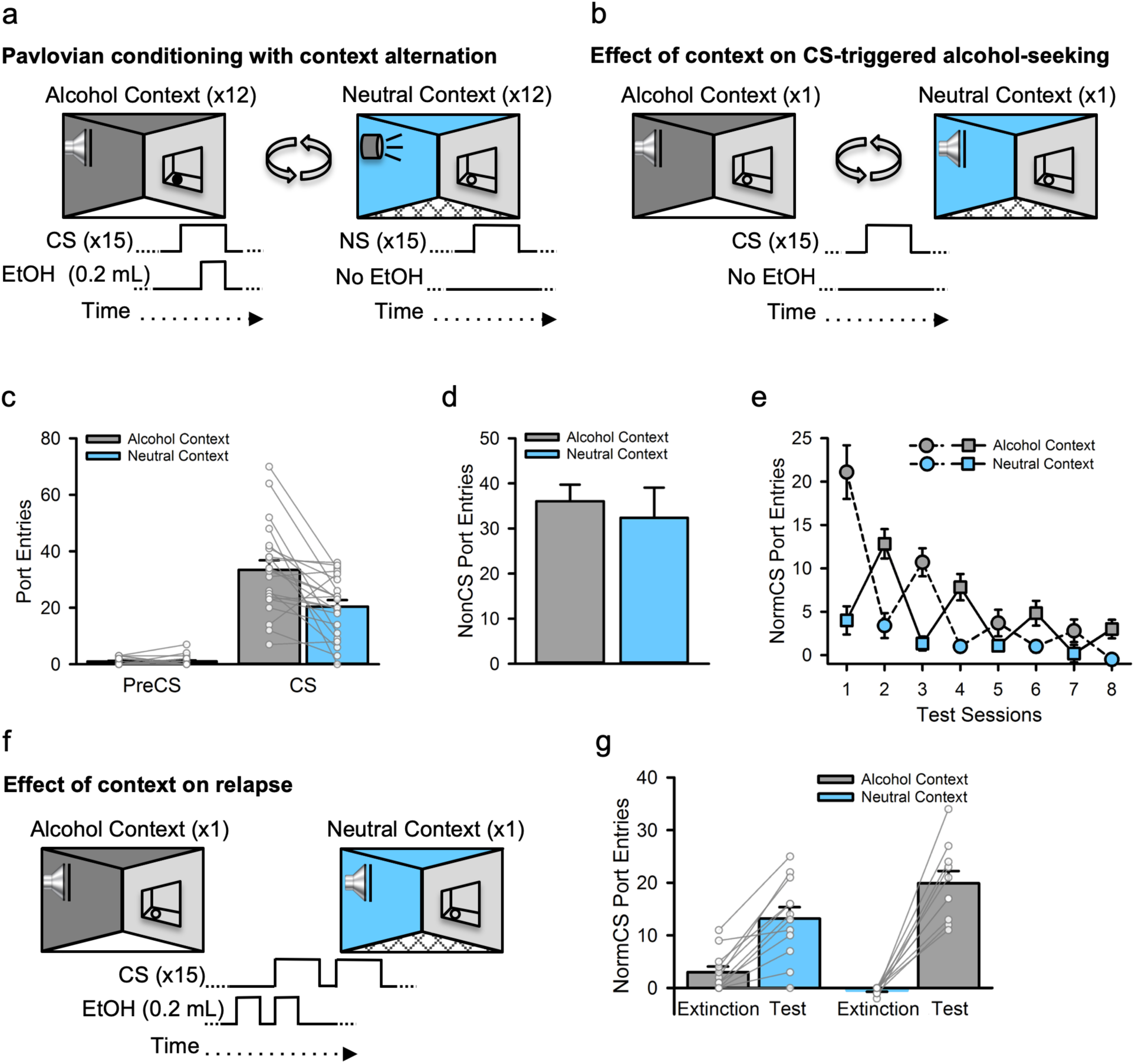
An alcohol context enduringly elevates alcohol-seeking triggered by a discrete cue and amplifies relapse. **a**, Rats (n=22) received Pavlovian conditioning sessions every other day in a distinct ‘alcohol context’ where a discrete auditory conditioned stimulus (CS) was paired with 15% ethanol (EtOH). On intervening days, rats were exposed to a different, ‘neutral context’ where a distinct, neutral auditory stimulus (NS) was presented without alcohol. **b**, CS-triggered alcohol-seeking was tested once each in the alcohol and neutral contexts by presenting the CS without alcohol. **c**, At test, CS port entries but not PreCS port entries were significantly elevated in the alcohol context compared to the neutral context [Context × Interval, *F*_(1, 20)_ =14.66, *p*=.001; CS port entries, *t*_(21)_ =3.736, *p*=.001], with no effect of context on **d**, port entries between CS trials [NonCS; Context, *F*_(1, 20)_ =.363, *p*=.554]. **e**, We then conducted 4 additional tests in the alcohol and neutral contexts to determine if the impact of the alcohol context on CS-triggered alcohol-seeking was a transient or enduring effect. Normalized CS port entries (i.e., CS minus PreCS port entries) waned across repeated tests in both contexts [Session, *F*_(3, 30)_ =18.435, *p*<.001], but exhibited a saw-tooth pattern that corresponded to elevated CS port entries in the alcohol context relative to the neutral context [Context, *F*_(1, 10)_ =30.671, *p*<.001; Context × Session, *F*_(3, 30)_ =5.885, *p*=.003]. Because of the counterbalanced experimental design, the first test occurred in the alcohol context (dashed line) for half the rats and in the neutral context (solid line) for the remainder. **f**, We then investigated the impact of context on relapse by presenting a drop of alcohol in the fluid port before and during the first CS presentation to reinstate extinguished CS port entries. Half of the rats received the reinstatement test in the neutral context following their last extinction session in the alcohol context and *vice versa* for the remainder. **g**, Relative to extinction, normalized CS port entries were reinstated in both contexts, but to a greater extent in the alcohol context than in the neutral context [Phase, *F*_(1,20)_ =108.887, *p*<.001; Phase × Reinstatement Context, *F*_(1, 20)_ =12.204, *p*=.002]. A Bonferroni-corrected post-hoc t-test indicated higher CS port entries at test in the alcohol context relative to the neutral context [*t*_(20)_ =2.112, *p*=.047]. All averaged data are shown as mean ± s.e.m. Data from individual rats are overlaid on the bar graphs.

Next, we examined CS-triggered alcohol-seeking in two tests, one that occurred in the alcohol context and one that occurred in the neutral context. At test, the CS was presented as in previous Pavlovian conditioning sessions, but no alcohol was delivered (**Fig. 1b**). CS port entries at test were higher than port-entries made during the PreCS interval, and elevated in the alcohol context relative to the neutral context (**Fig. 1c, S4a and d**). There was no impact of context on port entries made between CS trials (NonCS, **Fig. 1d**), the average latency to initiate a port entry following CS onset (**Fig. S4b**), or the duration of CS port entries (**Fig. S4c**). Thus, the alcohol context selectively elevated the number of CS port entries, without affecting other characteristics of the response form or general port-directed behaviour (NonCS).

To investigate the longevity of the context-induced elevation of CS port entries we conducted repeated tests in which the CS was presented without alcohol in the alcohol and neutral contexts on alternating days. The counterbalanced experimental design resulted in the first session of this phase occurring in the alcohol context for half of the rats and in the neutral context for the remainder. Normalized CS port entries (i.e., CS minus PreCS) decreased across test sessions but were elevated and maintained at a higher level for longer in the alcohol context (**Fig. 1e**). These findings reveal that CS-triggered alcohol-seeking was persistently elevated in the alcohol context.

Lastly, we examined the impact of context on relapse to CS-triggered alcohol-seeking produced by re-exposure to alcohol. The design for this test was counterbalanced so that rats that began extinction in the alcohol context received the relapse test in the neutral context, and *vice versa*. At test, 0.2 ml of 15% ethanol was delivered into the fluid port 90 s before the first inter-trial interval began and then again during the first of 15 CS trials (**Fig. 1f**). Normalized CS port entries increased significantly at test, compared to extinction. However, the difference between extinction and test was greater at test in the alcohol context, relative to at test in the neutral context (**Fig. 1g**). Thus, the alcohol context amplified the capacity of an orosensory alcohol prime to reinstate CS-triggered alcohol-seeking.

### Ventral tegmental area dopamine neurons are necessary for alcohol-seeking triggered by a discrete cue

Alcohol consumption, as well as exposure to contextual cues associated with operant alcohol self-administration, evokes dopamine release in the nucleus accumbens^13,14,12^ which is strongly innervated by dopaminergic neurons from the ventral tegmental area (VTA)^15^. Moreover, in a Pavlovian conditioning procedure in which a discrete CS predicted an aversive eye-puff, the firing rates of VTA dopamine neurons triggered by the CS were modulated by the context^16^. Here, we examined the role of the dopamine system and VTA dopamine neurons in alcohol-seeking triggered by a discrete cue for alcohol that was experienced in isolation from contextual alcohol cues.

First, we investigated the contribution of the dopamine system to CS-triggered alcohol-seeking by testing rats (n=11) in the neutral context following a systemic injection of saline, a dopamine D1-like (SCH23390, 10 μg/kg) or D2-like (eticlopride, 10 μg/kg) receptor antagonist (**Fig. 2a**). Pretreatment with either dopamine receptor antagonist reduced general port directed behaviour, as NonCS port entries were reduced relative to vehicle (**Fig. 2b, inset**). CS port entries were also significantly reduced by the D2 receptor antagonist eticlopride (*p*=.018), but not SCH23390 (*p*>1 after correction), relative to vehicle (**Fig. 2b**). Thus, dopamine neurotransmission at D1 and D2 receptors is needed for port directed behaviour that is not signaled by a discrete CS, while CS-triggered alcohol-seeking requires D2, but not D1 receptors. The decreases in both NonCS and CS port entries following eticlopride could suggest a non-specific locomotor deficit; however, we found previously that 10 μg/kg of eticlopride had no impact on CS port entries during a Pavlovian conditioning session in which the CS was paired with alcohol^17^.

**Fig 2.**
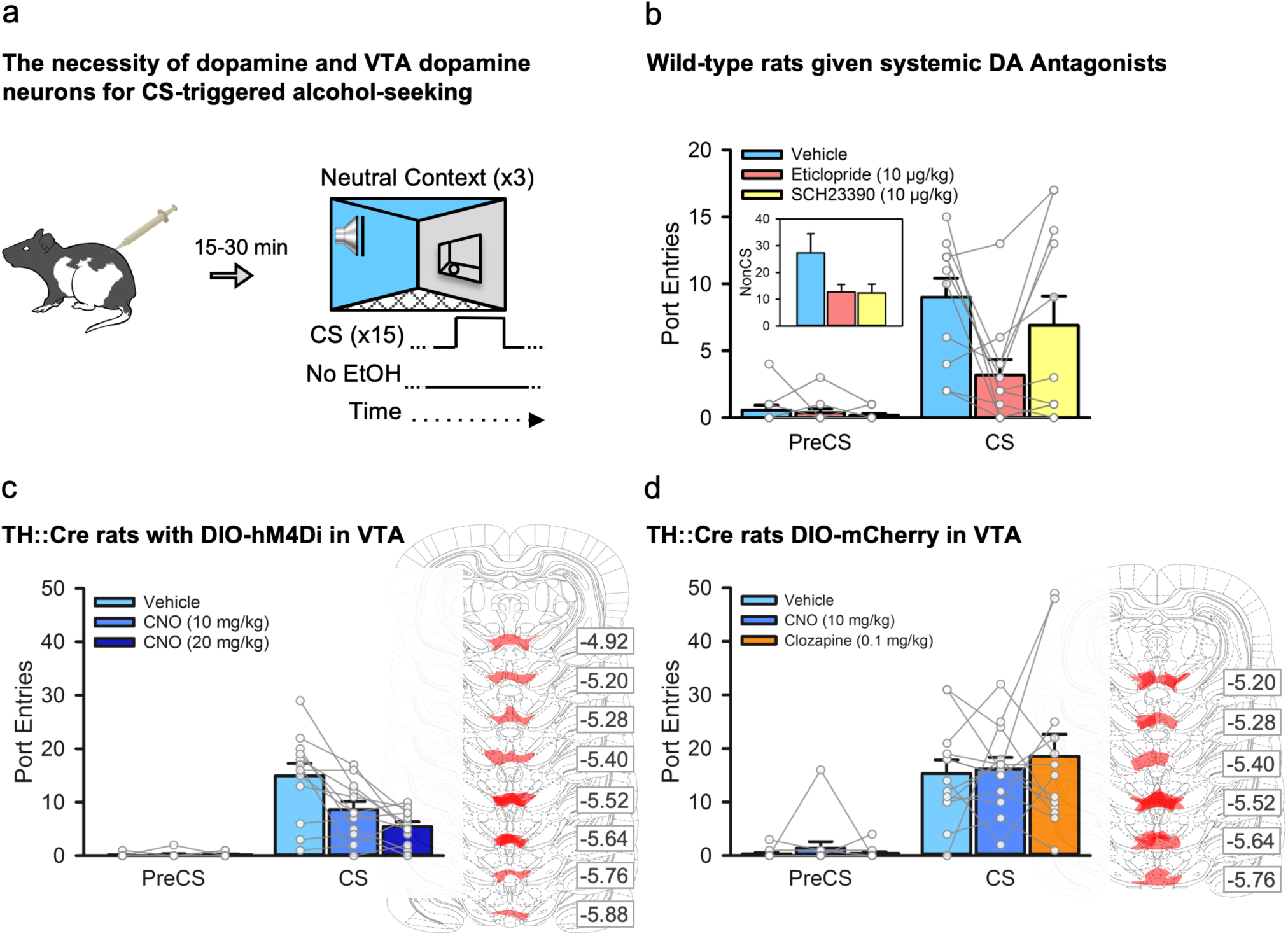
Ventral tegmental area dopamine neurons are necessary for alcohol-seeking triggered by a discrete cue. **a**, We examined the impact of inhibiting dopamine receptors, or inhibiting VTA dopamine neurons, on CS-triggered alcohol-seeking at test in the neutral context. **b**, In a subset (n=11) of the wild-type rats from figure 1, systemic administration of the dopamine D2-like receptor antagonist eticlopride [Interval × Treatment, *F*_(2, 20)_ =45.86, *p*=.028; Bonferroni-corrected *t*_(10)_ = 3.56, *p*=.018], but not the dopamine D1-like receptor antagonist SCH23390 (*p*>1 after correction), significantly reduced CS port entries relative to vehicle. NonCS port entries were also significantly reduced following dopamine receptor antagonist administration relative to vehicle [inset; Trea™ent, *F*_(2, 20)_ =4.17, *p*=.031]. **c**, Next, we used a chemogenetic approach in TH::Cre rats (n=12) that expressed the inhibitory designer receptor, hM4Di, in ventral tegmental area (VTA) dopamine neurons. At test, CS port entries were significantly reduced compared to vehicle [Interval by Trea™ent, *F*_(2, 22)_ =10.842, *p*=.001] following systemic administration of 10 mg/kg and 20 mg/kg of clozapine-n-oxide (CNO) (Bonferroni corrected *t*(11)=3.171, *p*=.03 for 10 mg/kg, and *t*_(11)_ =3.644, *p*=.012 for 20 mg/kg relative to vehicle). Maximal DIO-hSyn-hM4Di-mCherry expression in the VTA for each rat is shown in schematics from the atlas of Paxinos & Watson (2008). Numbers represent the anterior-posterior coordinate relative to bregma. **d**, The impact of CNO, and its parent compound clozapine, on CS-triggered alcohol-seeking was assessed in TH::Cre rats expressing the control DIO-mCherry construct in the VTA (n=13). Relative to vehicle, neither 10 mg/kg CNO nor 0.1 mg/kg clozapine significantly affected CS port entries [Interval × Trea™ent, *F*_(2, 24)_ =0.431, *p*=.655]. Maximal DIO-hSyn-mCherry expression for each rat is shown to the right. All averaged data are shown as mean ± s.e.m. Data from individual rats are overlaid on the bar graphs.

To circumvent the potential limitation of a general locomotor deficit produced by systemic administration of dopamine antagonists, and to verify a specific role for VTA dopamine neurons in CS-triggered alcohol-seeking we used a chemogenetic approach in TH::Cre rats to inhibit activity in a subset of VTA dopamine neurons. Twelve naïve TH::Cre male rats were microinfused bilaterally into the VTA with the double-floxed^18^ inhibitory designer receptor construct^19^ AAV8-hSyn-DIO-hM4Di-mCherry, resulting in the selective expression of the inhibitory designer receptor (hM4Di) in VTA dopamine neurons. This receptor inhibits firing of neurons when bound by the exogenous ligand clozapine-*n*-oxide (CNO)^19,20^. These rats then received twelve sessions of exposure to 15% ethanol in the home-cage (**Fig. S1b**) followed by Pavlovian conditioning with context alternation (**Fig. S2b and S3b**). Next, we tested CS-triggered alcohol-seeking by presenting the CS without alcohol in the neutral context 30 min after an intraperitoneal injection of vehicle or CNO (10 mg/kg or 20 mg/kg; **Fig. 2a**)^20,21^. PreCS responding was minimal at test, and CS port entries were reduced in a dose-related manner by CNO (**Fig. 2c**). The reduction in CS port entries could not be explained by general decreases in port directed behaviour, as NonCS port entries were similar across all tests (**Fig. S5a**). These data suggest that VTA dopamine neurons are critical for alcohol-seeking triggered by a discrete cue.

A naïve group of 13 TH::Cre rats received the same surgical, conditioning (**Fig S1c, Fig S2c and S3c**), and test procedures except that they were microinfused with a control virus construct (AAV8-hSyn-DIO-mCherry). At test, a within-subject design was used to treat rats with vehicle, CNO (10 mg/kg), or the parent compound of CNO, clozapine, at a dose (0.1 mg/kg) that CNO may have been reverse-metabolized to^22^. CS port entries at test in the neutral context were similar following vehicle, CNO, or clozapine (**Fig. 2d**), as were NonCS port entries (**Fig S5b**). Thus, the suppression of CS-triggered alcohol-seeking produced by chemogenetically inhibiting VTA dopamine neurons cannot be accounted for by non-specific effects of systemically injected CNO or its reverse-metabolism to clozapine.

Finally, we examined if silencing a similar population of VTA dopamine neurons would attenuate responding triggered by a CS that was associated with a natural reinforcer, sucrose (**Fig. S6a**). A naïve group of 18 TH::Cre rats was microinfused in the VTA with the AAV8-hSyn-DIO-hM4Di-mCherry and then given the same conditioning procedures as described earlier (**Fig S6b**) with the exception that CS presentations during Pavlovian conditioning sessions were paired with 10% sucrose solution instead of alcohol. In this study, sucrose-seeking was unaffected by CNO treatment as neither CS port entries (**Fig. S6c**) nor NonCS port entries (**Fig. S6c, inset**) differed across treatment conditions at test.

Together, these results show that chemogenetically inhibiting VTA dopamine neurons attenuated alcohol-seeking triggered by a discrete cue in a neutral context. This effect was not attributable to potential off-target effects of CNO or its parent compound clozapine, as we observed no effects of CNO or clozapine on CS port entries in TH::Cre rats that expressed a control fluorescent protein in the VTA. Interestingly, inhibiting VTA dopamine neurons using CNO in TH::Cre rats failed to reduce port entries triggered by a CS that predicted sucrose. This reinforcer selectivity may be due to the modest number of transfected neurons (∼25%; **Fig. 4c**), because the attenuation of feeding behaviour^23^ and hedonic orofacial responses to sucrose^24^ requires lesions that abolish at least ∼90% of total dopaminergic processes.

### Divergent mesolimbic dopamine circuits support alcohol-seeking triggered by discrete cues and contexts

VTA dopamine neurons project to the nucleus accumbens (NAc), and reports across multiple drug classes suggest dissociable roles for the ventral striatal core and shell subregions in drug-seeking triggered by discrete cues and contexts. Specifically, the NAc core appears necessary for drug-seeking triggered by discrete cues^12^, whereas the NAc shell appears to mediate alcohol-seeking and drug-seeking provoked by context^12,25^. This dichotomy is borne out by pharmacological research targeting the dopamine system. In the NAc core but not shell, drug-seeking triggered by a discrete cue is attenuated by microinfusion of a broad-spectrum dopamine receptor antagonist^26^ or a selective D1-like dopamine receptor antagonist^11^. Conversely, drug-seeking in context-based paradigms^11,26^ is reduced by these manipulations in the NAc shell, but not core. Based on this research, we hypothesized the VTA dopamine projections to the NAc core would be required for alcohol-seeking triggered by a discrete cue, whereas VTA dopamine projections to the NAc shell would be needed for the elevation of this behavior in an alcohol context.

We targeted dopaminergic projections from the VTA to the NAc core by microinfusing the AAV8-hSyn-DIO-hM4Di-mCherry construct into the VTA of 8 naïve TH::Cre rats and implanting bilateral cannulae targeting the NAc core. This approach enabled us to selectively silence dopaminergic VTA-to-NAc core projections by microinfusing CNO into the NAc core^20,27^. Following home-cage ethanol exposure (**Fig S1d**) and Pavlovian conditioning with context alternation (**Fig S2d and S3d**), we tested CS-triggered alcohol-seeking in both the alcohol context and the neutral context after CNO or vehicle microinfusions (**Fig. 3a**). As before, CS port entries were elevated over PreCS port entries and higher at test in the alcohol context, compared to the neutral context (**Fig. 3b**). CNO microinfusion into the NAc core reduced CS port entries in both the alcohol and neutral contexts (**Fig. 3b**). NonCS port entries were unaffected by CNO microinfusion or by context (**Fig. S5c**). Dopaminergic projections from the VTA to the NAc core were thus necessary for alcohol-seeking triggered by a discrete cue, irrespective of the context in which that discrete cue was experienced.

**Fig 3.**
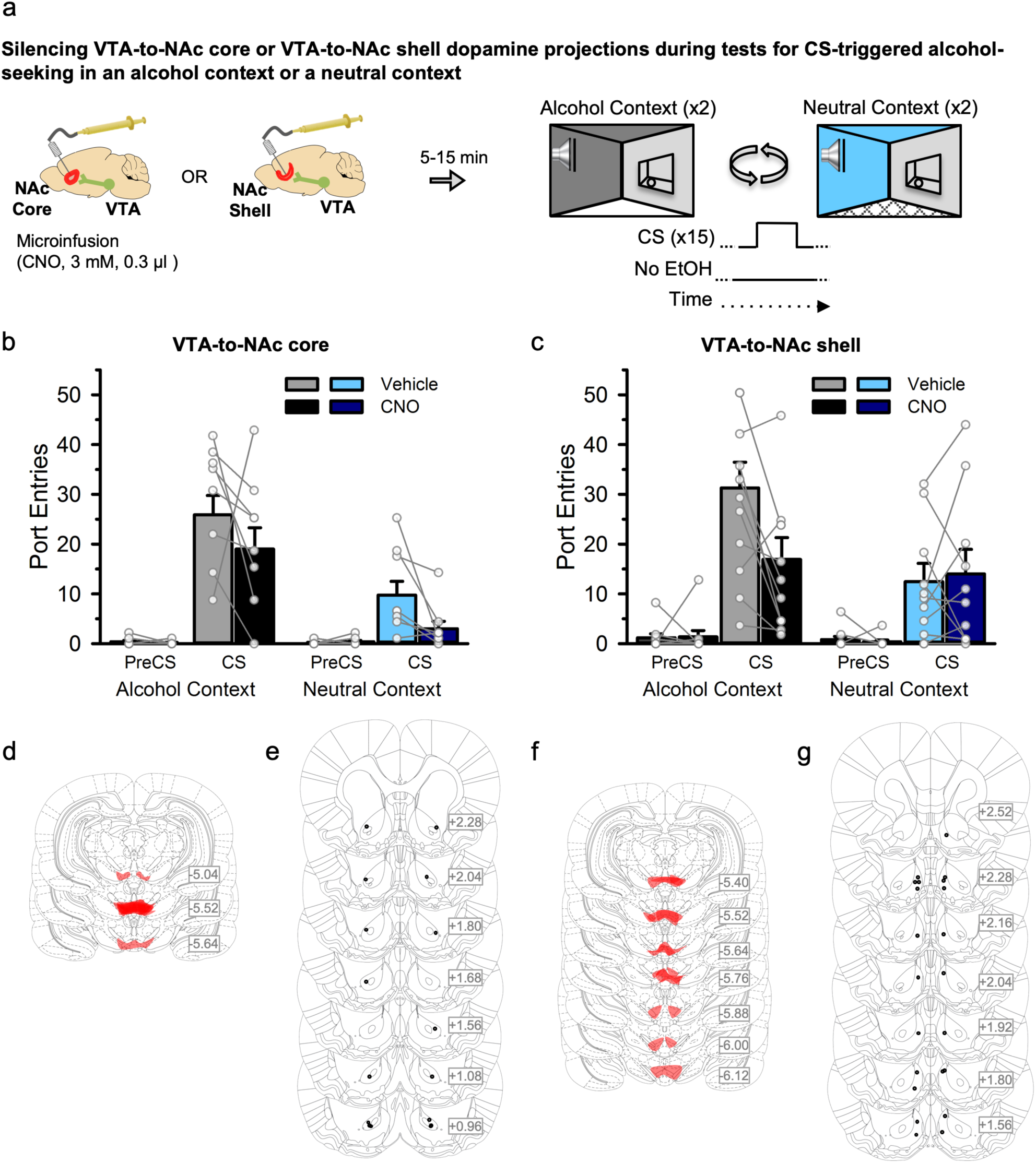
Divergent mesolimbic dopamine circuits support alcohol-seeking triggered by discrete cues and contexts. **a**, TH::Cre rats expressing the inhibitory designer receptor hM4Di in VTA dopamine neurons received Pavlovian conditioning with context alternation, and then CS-triggered alcohol-seeking was tested 4 times per rat, twice in the alcohol context and twice in the neutral context. Separate groups received a microinfusion of vehicle or CNO (3 mM, 0.3 μl) into either the NAc core (n=8) or shell (n=11) prior to each test. **b**, In rats with cannula targeting the NAc core, CS port entries were elevated at test in the alcohol context relative to the neutral context [Context by Interval, *F*_(1, 7)_ =44.840, *p*<.001]. Relative to vehicle, CS port entries were significantly reduced following CNO microinfusion in the NAc core in both contexts [Trea™ent × Interval, *F*_(1, 7)_ =5.763, *p*=.047; Context × Interval × Trea™ent, *F*_(1,7)_ =.002, *p*=.962]. **c**, In rats with cannula targeting the NAc shell, CS port entries were also elevated at test in the alcohol context relative to the neutral context [Context by Interval, *F*_(1, 10)_ =9.562, *p*=.011]. Relative to vehicle, there was no effect of CNO on CS port entries at test in the neutral context, but a significant reduction in CS port entries at test in the alcohol context [Context by Treatment by Interval, *F*_(1,11)_ =5.121, *p*=.047; CS port entries in neutral context *t*_(10)_ =-.367, *p*=.721; CS Port entries in alcohol context *t*_(10)_ =3.121, *p*=.011]. Histological results show mCherry expression in the VTA of the **d**, NAc core and **f**, NAc shell groups, as well as corresponding injector tip placements in the **e**, NAc core and **g**, NAc shell. All averaged data are shown as mean ± s.e.m. Data from individual rats are overlaid on the bar graphs.

Inactivating the NAc shell attenuates context-induced renewal of Pavlovian alcohol-seeking^12^ and microinfusing dopamine antagonists into the NAc shell attenuates context-induced renewal of drug-seeking^11^. We hypothesized that the elevation of CS-triggered alcohol-seeking in an alcohol context requires dopaminergic VTA-to-NAc shell projections. In a separate group of 11 naïve TH::Cre rats the same surgery, home-cage alcohol exposure (**Fig S1e**), conditioning (**Fig S2e and S3e**) and test procedures were performed as described for the VTA-to-NAc core experiment, with the exception that cannulae targeted the NAc shell (**Fig. 3a**). At test, CS port entries were elevated relative to PreCS port entries and higher in the alcohol context than in the neutral context (**Fig. 3c**). Relative to vehicle, microinfusions of CNO into the NAc shell reduced CS port entries, but this effect was selective for the alcohol context (*p*=.011; **Fig 3c**). The same manipulation had no effect on CS port entries at test in the neutral context (*p*=.721; **Fig 3c**). NonCS port entries were unaffected by CNO treatment, but were significantly lower in the neutral context than the alcohol context (**Fig S5d**). Thus, dopaminergic projections from the VTA to the nucleus accumbens shell are not required for alcohol-seeking triggered by a discrete cue, but support the enhancement of CS-triggered alcohol-seeking in an alcohol context. Together, these results suggest that a dopaminergic projection from the VTA to the NAc core subserves alcohol-seeking triggered by a discrete cue, while a divergent dopaminergic projection from the VTA to the NAc shell supports the elevation of this behavior in an alcohol context.

### Chemogenetic targeting of VTA dopamine neurons in TH::Cre rats, and modulation of synaptic input to medium spiny neurons in the nucleus accumbens core

The rat VTA contains an estimated 18,000 to 40,000 tyrosine hydroxylase positive neurons^15,28,29^ that make up 64 to 70%^15,29,30^ of the total cell population. We validated the selectivity of Cre for TH positive neurons, and the transfection efficiency of the AAV8-hSyn-DIO-hM4Di-mCherry construct in TH::Cre rats^18,19,31^. Brains from 4 TH::Cre^+/−^ rats that received home-cage alcohol exposure and Pavlovian conditioning with context alternation were used for immunohistochemical labeling of TH and amplification of the mCherry signal. In a single optical plane through the VTA at approximately bregma −5.5 mm (**Fig 4a and d**), 12.0 ± 1.5 cells were mCherry positive, 52.3 ± 6.3 were TH positive, and 11.6 ± 1.5 were colocalized (**Fig. 4b**). These counts produced an average selectivity of mCherry-expression for TH positive cells of 95.8% and a transfection efficiency of 24.8% (**Fig. 4c**), which are comparable to similar measures of selectivity in TH::Cre rats^20,31–35^. We also observed mCherry expressing TH positive processes in the NAc (**Fig. S7 and S8**), although this was not quantified.

**Fig 4.**
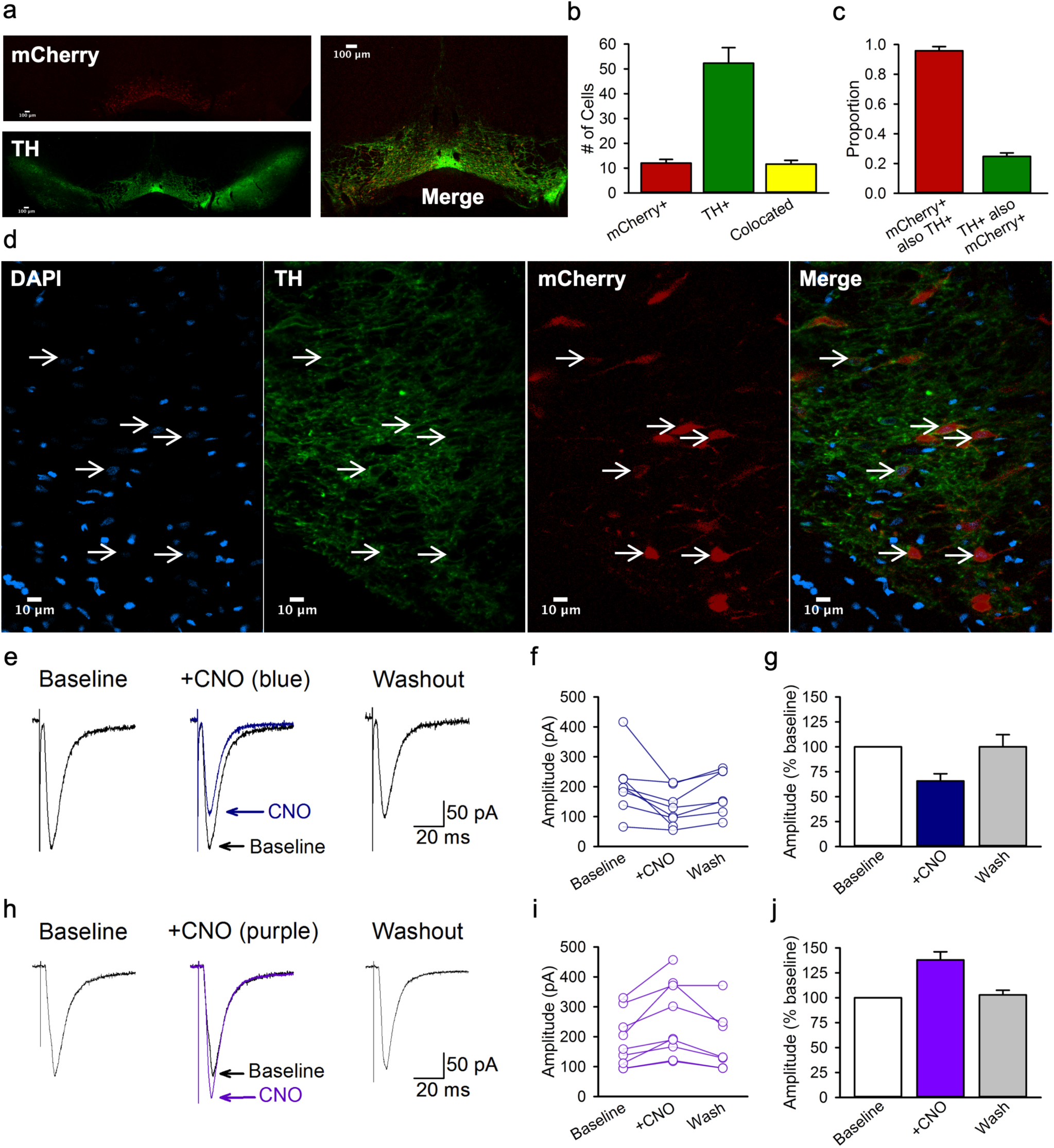
Chemogenetic targeting of VTA dopamine neurons in TH::Cre rats, and modulation of synaptic input to medium spiny neurons in the nucleus accumbens core. **a**, A representative coronal 4X optical plane through the VTA showing fluorescence indicative of mCherry (red; top; reporter for hM4Di) and TH (green; bottom), and a corresponding merged image at the right. **b**, The number of cells counted in each 300 × 300 μm optical section. **c**, Cell counts indicate the specificity of mCherry expression TH+ neurons as well as the proportion of TH+ neurons that were transfected. **d**, A representative area of a 40X image used for the analyses in **b** and **c**. Cells with colocalized TH and mCherry signals all have a nucleus indicated by DAPI (blue). **e**, Example traces show averaged excitatory post synaptic currents (EPSCs) for a NAc core medium spiny neuron before (baseline), during (+ CNO), and after (washout) 5 min application of 1 μM CNO to striatal slices containing hm4Di-expressing terminals from VTA dopamine neurons. **f**, Peak amplitudes of EPSCs recorded from individual medium spiny neurons innervated by hm4Di-expressing dopaminergic terminals. **g**, Normalized mean EPSC amplitudes for the same group of cells. CNO resulted in a marked reduction in EPSC amplitude to 65.7 ± 7.3% of baseline levels [*n*=8; *F*_(2, 13)_ =6.77, *p*=.01; *Newman-Keuls p*=.01]. Mean EPSC amplitude returned towards baseline values within 20 min of washout (93.9 ± 8.9 % of baseline). **h**, Example traces show averaged EPSCs for a medium spiny neuron before (baseline), during (+ CNO), and after (washout) 5 min application of CNO to striatal slices containing excitatory hM3Dq-expressing terminals from VTA dopamine neurons. **i**, Peak amplitudes of EPSCs recorded from individual medium spiny neurons innervated by hm3Dq-expressing terminals. **j**, Mean EPSC amplitudes for the same group of cells. CNO significantly increased EPSCs to approximately 137.9 ± 18.8% of baseline levels, [*n*=9, *F*_(2,14)_ =17.84, *p*<.001*; Newman-Keuls p*<.001]. Responses returned to baseline values within 20 min of washout (102.9 ± 4.7 % of baseline). All averaged data are shown as mean ± s.e.m.

We conducted *in vitro* intracellular recordings of excitatory synaptic potentials in medium spiny neurons (MSNs) in the NAc core to validate the ability of CNO to inhibit terminals of VTA dopamine neurons in the region where our *in vivo* manipulations occurred. Recordings were obtained within the NAc core, which showed strong mCherry fluorescence in the region of recorded neurons (**Fig. S8b and e**). Cells were identified as MSNs based on the characteristic slow, ramp-like depolarization of the membrane preceding spike firing^36,37^ and the characteristic action potential waveform followed by a prominent fast after hyperpolarization (**Fig. S8c**).

A 5-min bath application of 1 µM CNO to inhibit hM4Di-expressing terminals of VTA neurons within the NAc core did not produce significant changes in action potential number (*p*=0.28), width (*p*=0.14), or height (*p*=0.83) during injection of positive current steps. However, CNO significantly lowered action potential threshold from −43.2 ± 1.5 mV at baseline to −49.7 ± 2.4 in CNO (*p*=0.002), which is consistent with findings that dopamine *agonists* can shift threshold in the opposite direction, to more depolarized voltages^38^. CNO also resulted in a small decrease in the fast after hyperpolarization (5.7 ± .7 mV at baseline to 4.6 ± .7 in CNO, *p*=0.017). Application of CNO resulted in a marked reduction in excitatory post-synaptic current (EPSC) amplitude to 65.7 ± 7.3% of baseline levels (*p*=.01; **Fig. 4e-g**). In two neurons, subsequent application of the AMPA/NMDA receptor blocker kynurenic acid (50 μM) completely blocked the EPSC (data not shown), indicating that EPSCs were mediated by glutamatergic synaptic transmission. Thus, activation of inhibitory hM4Di designer receptors in the terminals of VTA neurons in the NAc core correspondingly inhibited excitatory postsynaptic responses in MSNs.

To confirm that the reduction in MSN EPSC amplitude was the result of CNO actions on designer receptors, we conducted a parallel analysis in NAc core slices that contained dopaminergic terminals expressing the excitatory designer receptor hM3Dq. We predicted that CNO acting on hM3Dq-expressing dopamine terminals in the NAc core would increase the amplitude of MSN EPSCs as dopamine has been reported to enhance MSN EPSCs in the NAc^39,40^. Accordingly, CNO significantly increased EPSCs to approximately 137.9 ± 18.8% of baseline levels, (*p*<.001; **Fig 4h-j**).

Thus, CNO acting on excitatory or inhibitory designer receptors expressed in dopaminergic terminals in the NAc core modulates the activity of post-synaptic MSNs. This bidirectional effect is consistent with at least three mechanisms by which dopamine influences the excitability of striatal MSNs. Firstly, dopamine can act synergistically at D1 and D2 receptors on MSNs to enhance MSN activity^41,42^. Secondly, glutamate co-release from VTA dopamine neurons can evoke EPSCs in post-synaptic MSNs that are insensitive to dopamine antagonists^43–45^. Thirdly, dopamine release from VTA terminals in the ventral striatum can act on corticostriatal terminals, which predominately express dopamine D1 relative to D2 receptors, to enhance glutamatergic input onto NAc MSNs^46–48^.

## Discussion

We report that alcohol-seeking triggered by a discrete cue was enduringly elevated, and reinstated to a higher level, in an alcohol context than in a familiar, neutral context. These findings highlight a role for context as a major determinant of relapse, particularly through interaction with discrete cues for alcohol. Alcohol-seeking triggered by a discrete cue was reduced by systemic administration of a dopamine D2 receptor antagonist and by chemogenetically silencing VTA dopamine neurons. Moreover, silencing dopaminergic projections from the VTA to the NAc core reduced CS-triggered alcohol-seeking irrespective of context, whereas silencing VTA projections to the NAc shell only reduced the elevation of this behaviour in an alcohol context. These data ascribe unique roles to divergent mesolimbic dopamine circuits in alcohol-seeking triggered by a discrete cue and the enhancement of CS-triggered alcohol-seeking in a context associated with alcohol.

The rat ventral tegmental area contains a large population of dopamine cell bodies that project to diverse striatal and prefrontal targets^15,35,49^. Dopamine cell bodies are organized along a medio-lateral gradient in the VTA, with cells originating near the midline projecting predominantly to medial areas of the striatum, including the medial NAc shell, and more lateral areas projecting to the NAc core^35,50,51^. Coupling cre-dependent designer receptor constructs with transgenic TH::Cre rats allowed for highly specific targeting of VTA dopamine neurons. We observed designer receptor expression encompassing the anterior-posterior and medio-lateral expanse of VTA dopamine neurons that project to both the NAc core and shell, with relatively minimal expression in the substantia nigra.

Although VTA dopamine neurons are strongly implicated in learning *via* prediction error ^32,52,53^, no study has directly examined their contribution to CS-triggered alcohol-seeking. We found that systemic blockade of D2 dopamine receptors or chemogenetic silencing of VTA dopamine neurons reduced alcohol-seeking triggered by a discrete cue in a neutral context. These results are the first to demonstrate the necessity of VTA dopamine neurons in alcohol-seeking triggered by a discrete cue, and are consistent with studies suggesting a role for cholinergic^54^ and glutamatergic^55^ signaling within the VTA in cue conditioning with alcohol. Interestingly, the lower dose of CNO that effectively reduced CS-triggered alcohol-seeking had no effect on CS-triggered sucrose-seeking. This result may be explained by high redundancy in the neural circuitries that support responding for natural food reinforcers^23,24^. Our silencing of a modest proportion of VTA dopamine neurons may have been sufficient to disrupt CS-triggered alcohol-seeking but not sucrose-seeking.

There is a longstanding hypothesis based on lesion and pharmacological data that the NAc core, but not NAc shell, mediates responding triggered by discrete Pavlovian cues in a dopamine-dependent manner^11,12,56–59^. Supporting this, observational studies reveal the development of dopamine transients in the NAc core that are time locked to the onset and offset of food-predictive cues^60^, and that track closely with the earliest reliable predictor of reinforcement^61^. Remarkably, pairing a cue with optogenetic stimulation of VTA dopamine neurons or their projections to the NAc core, but not the NAc shell, leads to the cue acquiring incentive motivational properties^35^. This decisive finding underscores a central role for a VTA-to-NAc core dopamine pathway in Pavlovian learning about discrete cues^35^. Studies in which alcohol is used as the reinforcer also implicate dopamine neurotransmission within the NAc core in the expression of alcohol-seeking behaviour^12,17,62,63^. In the present research, after carefully controlling for the influence of context on behavioural responding, we provide evidence for a critical role of dopamine neurons projecting from the VTA to the NAc core in the expression of alcohol-seeking triggered by a discrete cue. The exact nature of the involvement of this circuit in alcohol-seeking triggered by a discrete CS might arise from the NAc core being necessary for orchestrating behavior in response to the best predictors of reinforcement.

Silencing VTA dopamine inputs to the NAc shell had no effect on CS-triggered alcohol-seeking in a neutral context, but selectively reduced the elevation of this behaviour in an alcohol context. This cross-context comparison supports the conclusion that the dopamine projection from the VTA to the NAc shell is not necessary for responding to the CS *per se*, but rather mediates the capacity of the alcohol context to augment CS-triggered alcohol-seeking. This finding corroborates an emerging hypothesis that the NAc shell is an important hub in neural circuits that mediate the impact of drug-contexts on drug-seeking behaviour^12,58,64–66^. Interestingly, optogenetic inhibition of medium spiny neurons that project from the NAc shell onto GABA interneurons in the VTA, causing a net inhibition of VTA dopamine neurons, attenuated context-induced renewal of alcohol-seeking in an operant conditioning procedure^64^. We found that chemogenetically silencing dopamine terminals in the NAc core reduced the strength of synaptic inputs onto MSNs. A similar reduction in inputs to the NAc shell may have reduced the excitability of MSNs that project back onto GABA interneurons in the VTA^51,64,67^, contributing to inhibition within the VTA that may regulate the impact of drug contexts on drug-seeking behaviour.

The selective reduction in CS-triggered alcohol-seeking in the alcohol context produced by silencing VTA dopamine inputs to the NAc shell substantiates earlier findings using a similar procedure, in which pharmacological inactivation of the NAc shell attenuated CS-triggered alcohol-seeking in an alcohol context but not in a neutral context^5^. However, this global inhibition of the NAc shell also increased port entries during the NonCS interval^5^. Alcohol-seeking in response to a discrete CS versus general port-directed behaviour may be mediated by separate inputs to the NAc shell that require a circuit-specific approach to be delineated.

Altogether, the present data support the hypothesis that the behavioural regulation of alcohol-seeking by discrete cues and contexts is controlled by divergent but potentially overlapping neural circuits^68,69,12^. Whether the separable roles of mesolimbic dopamine projections in this behaviour arise from circuit architecture in the VTA^35^ or the target regions^70^ is unknown. Nonetheless, elucidating the function of separable mesolimbic dopamine circuits provides insight into the understanding and prevention of cue-triggered alcohol-seeking and relapse.

In people with alcohol use disorder, environmental cues that predict alcohol can trigger reactivity that promotes alcohol-seeking and facilitates relapse^2,71^. Extinguishing these reactions by presenting patients with cues in the absence of alcohol has been used for treatment, but this form of extinction therapy has only limited efficacy. We show that alcohol-seeking triggered by a non-extinguished CS is elevated in an alcohol context, and that the extinction of alcohol-seeking is prolonged in an alcohol context relative to a neutral context. Furthermore, exposure to an oral alcohol prime after extinction reinstated alcohol-seeking in both contexts, but to a greater level in the alcohol context. These results are consistent with the capacity of alcohol cues to trigger relapse outside the treatment facility following extinction training, and suggest that patients may be particularly vulnerable to relapse when confronted with discrete cues for alcohol in an alcohol context. In an additional analysis we show that elevated CS-triggered alcohol-seeking at test is better predicted by the rate of learning during acquisition than by other measures of port-directed behaviour or alcohol consumption (**Fig S4e**). These results suggest that the alcohol-seeking we observe at test reflects a behaviour that is shaped throughout acquisition to a greater extent than it reflects a natural proclivity to drink alcohol or respond to alcohol-associated cues. It may also be true in humans that the susceptibility to cue- and context-induced relapse depends on the extent to which a person experiences repeated pairing of cues, contexts, and alcohol. Understanding the psychological and neural processes that govern the independent and combined effects of discrete and contextual alcohol cues on alcohol-seeking behaviour is essential for devising new, efficacious strategies that will prevent cues from instigating relapse.

In conclusion, we report a robust and enduring impact of alcohol contexts on alcohol-seeking triggered by a discrete cue that has direct implications for people with alcohol use disorder. Through systematic manipulation of context in our experimental design, we found that divergent mesolimbic dopamine circuits were engaged when a discrete cue for alcohol was experienced in a context where alcohol was previously consumed, relative to a familiar, neutral context. Specifically, we identify a point of separability in this circuitry at the level of ventral striatal targets for VTA dopamine neurons, highlighting that target region is on par with neurotransmitter content or anatomical origin as a means to delineate unique functional roles for distinct neural circuits. However, to fully understand the neural substrates that control problematic alcohol-seeking will likely require the ability to control the activity of circuitries that differ across even more subtle characteristics, such as their genetic profile^72^, or protein expression levels^73,74^.Ultimately, refining our molecular and anatomical understanding of the neural substrates that control alcohol-seeking triggered by discrete and contextual cues, as well as their interaction, may allow for the development of more precise interventions to prevent alcohol-seeking and relapse in humans.

## Methods

### Apparatus

Behavioral training and testing was conducted using equipment and software from Med-Associates Inc. (St. Albans, VT, USA). There were 12 conditioning chambers (ENV-009A), each contained within a fan-ventilated (84-88 dB), sound-attenuating, melamine cubicle (53.6 × 68.2 × 62.8 cm; built in-house). Each chamber had a stainless-steel bar floor (ENV-009A-GF), paneled aluminum sidewalls, and a clear polycarbonate rear wall, ceiling and front door. The right wall featured a fluid port (17.5 cm from the rear wall and 9 cm from the front door) that contained two circular fluid wells (ENV-200R3AM). Fluid delivery into one well occurred through a 20 ml syringe attached to a pump (PHM-100, 3.33 rpm) located outside the cubicle. Entries into the fluid port were measured by interruptions of an infrared beam (ENV-205M) across its entrance, and recorded to a computer using Med PC-IV software which also controlled fluid delivery and stimulus presentations. The upper left wall of the conditioning chamber featured a clicker stimulus (ENV-135M, 5 Hz, 8 dB above background) and a continuous white noise stimulus generator (ENV-225SM, 8 dB above background), as well as a white house-light (ENV-215M).

### Solutions and Reagents

Odours used to differentiate contexts were prepared by adding lemon oil (SAFC Supply Solutions, St-Louis, MO, USA; #W262528) or benzaldehyde (almond odor; OMEGA Chemical Company Inc., Levis, QC, Canada; #B37-50) to tap water to produce separate solutions of 10% concentration (v/v). Ethanol (15 %, v/v) was prepared fresh weekly by diluting 95% ethanol in tap water and was stored and used at room temperature. The D2-like receptor antagonist eticlopride (C_17_H_25_ClN_2_O_3_. HCl Sigma Aldrich #E101) and the D1-like receptor antagonist SCH23390 (C_17_H_18_ClNO. HCl Sigma Aldrich #D054) were dissolved in sterile 0.9% saline to make 10 μg/ml concentrations of each solution. CNO (Clozapine-*n*-oxide; Tocris #4936; NIMH C-929) for systemic administration was dissolved in 5% dimethyl sulfoxide and 95% sterile 0.9% saline to make 10 or 20 mg/ml CNO concentrations. Clozapine for systemic administration (AdooQ #A10236-500) was dissolved in 5% dimethyl sulfoxide and 95% sterile saline to make a 0.1 mg/ml solution. CNO (abcam #ab141704) was dissolved in sterile 0.9% saline at a concentration of 0.3 mM for intracerebral microinfusions, and was dissolved in artificial cerebrospinal fluid at a concentration of 1 μM for bath application during *in vitro* electrophysiology experiments. Viral vectors were supplied by the University of North Carolina Vector Core [AAV8-hSyn-DIO-hM4D(Gi)-mCherry (titer 5.3 or 4.6×10^12^), AAV8-hSyn-DIO-hM3D(Gq)-mCherry (titer 5.9×10^12^)] or Addgene [AAV8-hSyn-DIO-hM4D(Gi)-mCherry (titer 2.06×10^12^), AAV8-hSyn-DIO-mCherry (titer 2.1×10^13^)].

### Subjects

Twenty-four male, Long-Evans (220-275 g on arrival; INVIGO, previously Harlan) and 94 outbred male, Long-Evans heterozygous TH::Cre^+/−^ rats (bred in-house) were single-housed on a 12 h light-dark cycle at 21±2°C at approximately 40-50% humidity with unrestricted access to standard rat chow (Charles River Rodent Diet #5075) and water. All rats had access to a nylabone^™^ chew toy for enrichment. Founder TH::Cre rats were kindly provided by Dr. Karl Deisseroth^31^. All procedures were conducted in the light phase. All experimental procedures were approved by the Animal Research Ethics Committee at Concordia University and complied with regulations provided by the Canadian Council on Animal Care.

In total, 11 rats failed to acquire Pavlovian conditioning with ethanol as the unconditioned stimulus (an average of <5 CS port entries per session on the last two sessions of training), 6 rats had missed cannula placements, 2 rats experienced a programming error during conditioning and, 1 rat had the house-light burn out during a test session. Data from these rats were excluded.

### Surgery

Rats were anesthetized in an induction chamber with 5% isoflurane in oxygen and maintained at 2-3% isoflurane during surgery. Immediately after, the skull was shaved to expose the scalp, subjects were secured in the stereotaxic frame and then administered 0.1 ml/kg of atropine subcutaneously (s.c.) to prevent respiratory congestion, Proviodine^™^ was used to disinfect the incision site and tear-gel^™^ was applied to the eyes. Bilateral ventral tegmental area (VTA) microinfusions of 1 μl of AAV8-hSyn-DIO-hM4D(Gi)-mCherry, AAV8-hSyn-DIO-mCherry, or AAV8-hSyn-DIO-hM3D(Gq)-mCherry viral vectors were made at 0.1 μl/min over 10 min through a 26 gauge injector connected with PE20 tubing to a Hamilton microinjection syringe secured to a Harvard Apparatus, Pump 11 Elite. VTA virus infusions were delivered at a 10° angle using the following coordinates (in mm) from bregma: AP −5.5, ML ±1.84, DV −8.33. Following the 10 min infusion, the injector was left in place for 10 min for diffusion. 26-gauge guide cannulae (PlasticsOne C315G-SPC) were then implanted bilaterally to target the nucleus accumbens (NAc) core or shell. Cannulae were implanted 3 mm dorsal to the injection site using the following coordinates (in mm) at a 10° angle: NAc core AP +1.2, ML +/- 3.23, DV - 7.11, and NAc shell AP +1.68, ML +/- 2.23, −7.35. After surgery, rats were administered the analgesic buprenorphine (0.1 mg/kg, s.c.). Antiseptic (polysporin^™^) and a local analgesic (lidocaine^™^) were applied to the skin surrounding the incision. Rats recovered from anesthetic in a clean cage that was warmed by a heating lamp. Standard rat chow was mixed with sucrose and water and placed in the home cage. Rats were given at least one week to recover from surgery, during which time their overall health and body weight were monitored daily.

### Behavioural procedures

#### Exposure to alcohol in the home-cage

Intermittent access to ethanol solutions produce increases in consumption as rats become familiar with the taste and pharmacological effects of ethanol^17,75^–^80^. Every other day rats were given 24 h access to both a 15% ethanol solution and to tap water in separate containers in the home-cage. Ethanol was provided in a 100 ml graduated cylinder fitted with a rubber stopper containing a sipper tube with a metal ball bearing to minimize spillage. Water was provided in a standard 500 ml water bottle with the same sipper tube. On the intervening days the ethanol cylinder was removed but the water bottle was left in place. Ethanol cylinders and water bottles were placed on opposite sides of a standard cage lid, and the position was alternated across ethanol access sessions to control for side-preferences. Containers were weighed before and after every 24 h session. Filled ethanol cylinders and water bottles were placed onto two empty cages and handled in the same manner, and the amount of fluid lost from these containers was used to estimate spillage. Rats that consumed less than 1 g/kg of ethanol on the 6^th^ session were given 2 sessions of access to 5% ethanol and then returned to 15% ethanol for the remaining sessions.

In experiment 4 which used 10% sucrose instead of ethanol, rats were exposed to 10% sucrose in the home-cage for two, consecutive 24 h periods and consumption was monitored as described above.

#### Pavlovian conditioning with context alternation

Rats were first habituated to transport and the conditioning chambers. On the last day of home-cage alcohol exposure, rats were brought to the behavioural testing room in their home-cages and individually handled. On the two subsequent days, rats were habituated to the conditioning chambers with Context 1 on the first day, and to Context 2 24 h later. Context 1 included black walls, a clear polycarbonate floor, and a lemon odour. Context 2 included clear Plexiglas walls, a wire-mesh floor, and an almond odour. Odours were applied to a petri dish placed in the waste-pan located underneath the chamber floor. Habituation sessions began with illumination of the house-light after 1 min, and lasted 20 min. Entries into the fluid port were recorded in each session. Sessions for habituation to the room and to the conditioning chambers occurred at approximately the same time of day as subsequent training and test sessions.

Next, rats underwent Pavlovian conditioning with context alternation to pair a discrete auditory CS with ethanol delivery into the fluid port in a distinct alcohol context. On the basis of ethanol consumption (g/kg) in the home-cage, rats were assigned to one of two groups that consumed similar amounts of ethanol overall. These two groups received twelve Pavlovian conditioning sessions (73.5 min duration) in either Context 1 or Context 2 every other day, and exposure to a neutral context (see below) on alternating days. Sessions began with a two-minute delay during which port entries were recorded. After the delay, the house-light was illuminated and the first inter-trial interval began. In each session, 15 trials of an auditory CS (10 s; continuous white noise or clicker 5 Hz) occurred on a variable-time (VT) 260 s schedule, with inter-trial intervals (ITIs) as the time between CS offset and the next CS onset of 140, 260, or 380 s. With every CS presentation, 0.2 ml of 15% ethanol was dispensed into the fluid port over the last 6 s of the CS, resulting in a total of 3 ml of ethanol delivered per session. Ports were checked at the end of each session to make sure that all the ethanol was consumed.

On the days intervening the Pavlovian conditioning sessions all rats were exposed to a distinct ‘neutral context’ for 73.5 min sessions. The neutral context was the same conditioning chamber wherein Pavlovian conditioning occurred but was configured as a different context type. For example, rats assigned to context 1 as the alcohol context received context 2 as the neutral context, and *vice versa*. Half of the rats in each group were exposed to the alcohol context in the first session, and half of the rats were exposed to the neutral context in the first session. In neutral context exposure sessions, the house-light was illuminated after a two-minute delay and an auditory neutral stimulus (NS; clicker or white noise) that was different from the assigned CS was presented on a 260 s VT schedule, but no ethanol was delivered in these sessions. Syringe pumps were activated on a similar schedule as during Pavlovian conditioning sessions, but syringes were not placed on the pump.

This protocol of Pavlovian conditioning with context alternation was conducted identically in all experiments, with 2 exceptions. First, in our initial behavioural study (experiment 1), we compared the effect of having an NS present or absent from the neutral context. We found no difference in acquisition or subsequent test results in rats that had or had not received an NS in the neutral context, and all subsequent studies included an NS in the neutral context. Second, in experiment 4 in which we examined the impact of silencing VTA dopamine neurons using chemogenetics on CS-triggered sucrose seeking, 10% sucrose was used as the unconditioned stimulus.

#### Tests for CS-triggered alcohol-seeking

Following the training procedure described above, tests were conducted as specified below for each experiment. All tests occurred 24 h after a training session. In all tests, the CS was presented on the same VT 260 s schedule as in prior Pavlovian conditioning sessions, but without ethanol delivery. Syringe pumps were activated but no ethanol was delivered. The order of the test context and all treatment conditions were fully-counterbalanced in all experiments.

#### Experiment 1: The effect of context on CS-triggered alcohol seeking and reinstatement

Using the method described above, we tested CS-triggered alcohol-seeking in the alcohol context for half the rats and in the neutral context for the remainder (n=22). Rats then had two re-training sessions, one per day in each context as during training, prior to a second test for CS-triggered alcohol-seeking conducted the following day in the opposite context from that used in test 1.

At 24 h after the last test, we conducted repeated daily test sessions to assess the endurance of CS-triggered alcohol-seeking in the alcohol and neutral contexts. All rats were exposed to the CS, without ethanol, in both the alcohol and neutral contexts on alternating days for a total of eight sessions (four in each context). For half the rats, the first test in the series of repeated tests occurred in the alcohol context, and for the remainder it occurred in the neutral context. Test sessions alternated daily between the 2 contexts, and in each test the CS was presented on a VT 260 s schedule, but ethanol was withheld. Syringe pumps were activated as during training.

After the last repeated test session, in which CS-triggered alcohol-seeking had extinguished to similar levels in both contexts, we examined the impact of the alcohol context on CS-triggered alcohol-seeking in a model of relapse. All rats received a reinstatement test in the opposite context from the one in which the last of the repeated test sessions occurred. Thus, half of the rats received a reinstatement test in the alcohol context (n=10) following a session in the neutral context and the other half received a reinstatement test in the neutral context (n=12) following a session in the alcohol context. At test, a 0.2 ml drop of ethanol was dispensed over 6 s into the fluid port 30 s after rats were placed in the chamber. A second 0.2 ml drop of ethanol was delivered during the last 6 s of the first CS presentation, with no more ethanol delivered for the remainder of the test session. The purpose of this ethanol prime was to serve as a reminder of the taste and smell of ethanol.

#### Experiment 1b: Role of dopamine in CS-triggered alcohol-seeking in a neutral context

A subset of rats from experiment 1 (n=11) received 3 Pavlovian conditioning sessions intervened by 3 sessions of exposure to the NS in the neutral context. All rats received two 1 ml/kg s.c. saline habituation injections, 15 min before each of the last two training sessions, one in each context. Next, three tests for CS-triggered alcohol-seeking in the neutral context were conducted after systemic s.c. administration of vehicle, the dopamine D2 receptor antagonist eticlopride (10 μg/kg), or the D1 receptor antagonist SCH23390 (10 μg/kg)^17,81^. Test sessions were alternated with single Pavlovian conditioning sessions in the alcohol context. Test injections occurred 15 min before test sessions began.

#### Experiment 2: Effect of chemogenetically silencing VTA dopamine neurons on CS-triggered alcohol-seeking in the neutral context

We tested the necessity of VTA dopamine neurons for CS-triggered alcohol-seeking in a neutral context. TH::Cre rats (n=12) received two 1 ml/kg saline habituation injections intraperitoneally (i.p.), one in each context, 30 min before each of the last two training sessions. Next, three tests for CS-triggered alcohol-seeking in the neutral context were conducted after systemic i.p. administration of vehicle, CNO at 10 mg/kg, or CNO at 20 mg/kg^20,21^. Tests were intervened by four retraining sessions, two in the alcohol context and two in the neutral context.

#### Experiment 3: Effect of potential off-target effects of CNO and its parent compound clozapine on CS-triggered alcohol-seeking in the neutral context

We tested the potential off-target effects of CNO and its parent compound, clozapine, on CS-triggered alcohol-seeking in a neutral context. TH::Cre rats (n=13) received two 1 ml/kg saline habituation injections (i.p.), one in each context, 30 min before each of the last two training sessions. Next, three tests for CS-triggered alcohol-seeking in the neutral context were conducted 30 min after systemic i.p. administration of vehicle, 10 mg/kg CNO, or 0.1 mg/kg clozapine. Tests were intervened by two retraining sessions, one in the neutral context and one in the alcohol context.

#### Experiment 4: Effect of chemogenetically silencing VTA dopamine neurons on CS-triggered sucrose-seeking in the neutral context

We tested the necessity of VTA dopamine neurons for CS-triggered sucrose-seeking in a neutral context. TH::Cre rats (n=18) received two 1 ml/kg saline habituation injections (i.p.), one in each context, 30 min before each of the last two training sessions. Two tests for CS-triggered alcohol-seeking in the neutral context were then conducted in the neutral context 30 min after systemic i.p. administration of vehicle or 10 mg/kg CNO, according to a counterbalanced, within-subject design. Between tests, rats received three consecutive retraining sessions, two in the sucrose context intervened by one in the neutral context.

#### Experiment 5: Effect of chemogenetically silencing the dopaminergic projection from the VTA to the NAc core on CS-triggered alcohol-seeking in the alcohol and neutral contexts

We tested the necessity of the VTA to NAc core projection for CS-triggered alcohol-seeking in an alcohol and neutral context. TH::Cre rats (n=8) received a saline microinfusion (0.15 μl over 1 min) into the NAc core 5-15 min before a training session in the alcohol and neutral contexts to habituate them to this process. After the last training session, the first of four tests for CS-triggered alcohol-seeking was conducted, in which a similar number of rats were tested in their alcohol or neutral context after receiving a vehicle or CNO (3 mM) microinfusion (0.3 μl over 1 min) 5-15 min before the test session^20,27,82^. Between each test, rats received two retraining sessions, one in the neutral context and one in the alcohol context.

#### Experiment 6: Effect of chemogenetically silencing the dopaminergic projection from the VTA to the NAc shell on CS-triggered alcohol-seeking in the alcohol and neutral contexts

We tested the necessity of the VTA to NAc shell projection for CS-triggered alcohol-seeking in an alcohol and neutral context. TH::Cre rats (n=11) received a saline microinfusion (0.15 μl over 1 min) into the NAc shell before a training session in the alcohol and neutral context to habituate them to this process. After the last training session, the first of four tests for CS-triggered alcohol-seeking was conducted, in which a similar number of rats were tested in their alcohol or neutral context after receiving a vehicle or CNO (3 mM) microinfusion (0.3 μl over 1 min) 5-15 min before the test session. Between each test, rats received two retraining sessions, one in the neutral context and one in the alcohol context.

#### Experiment 7: Immunohistochemistry for mCherry to visualize hM4Di and tyrosine hydroxylase in TH+ neurons

We examined the selectivity of designer receptors to be expressed in TH+ neurons and the transfection efficiency of our viral constructs in TH::Cre rats (n=4). Rats were deeply anesthetized with euthanyl^™^ (sodium pentabarbitol 240 mg/kg i. p.) and perfused with approximately 250 ml of 0.02 M phosphate buffered saline (PBS; pH 7.2), and 150 ml of 4% paraformaldehyde in 0.02 M PBS (pH 7.2). Immediately following perfusion, brains were removed and stored in 50 ml of a 4% paraformaldehyde 30% sucrose solution for cryoprotection for approximately 2-3 days. They were then removed from the sucrose solution and stored at − 80°C until they were sectioned (40 μm thick) using a cryostat at −20°C and thaw-mounted. Slides were removed from the −20°C freezer and allowed to dry, covered by a lid, in a fumehood overnight. The entire slide, excluding the labeled portion, was then outlined with an ImmEdge^™^ hydrophobic barrier pen (Vector labs #H-4000) and then washed twice by pipetting 500 μl of 0.01 M PBS onto the slide for 1 minute. After the second wash, 500 μl of 10% normal donkey serum (NDS; Sigma-Aldrich #D9663) in 0.01 M PBS plus 0.3% Triton X-100 (PBST) was applied to the slide for 30 min. After a 1 min wash in 0.01 M PBS, 500 μl of mouse anti-mCherry (1:1000; abcam #ab125096) and rabbit anti-TH (1:100; EMD Millipore #ab152) in 10% NDS PBST was applied to the slides and left to incubate for 48 h at room temperature. Slides were then washed 3 times with 500 μl of 0.01 M PBS for 5 min. Then, 500 μl of donkey anti-mouse IgG (H+L) alexa 594 (1:200; Jackson ImmunoResearch labs #715-585-150) and donkey anti-rabbit IgG (H+L) alexa 488 (1:200; Jackson ImmunoResearch labs #711-545-152) in 0.01 M PBS was applied to the slides and left to incubate for 24 h at room temperature. After the secondary incubation, slides were left to dry covered at room temperature for 2 h and then coverslipped with vectasheild^™^ (Vector labs #H-1200) and imaged immediately or stored at 4°C until imaging.

We performed colocalization analyses to verify that our designer receptor viral constructs were selectively expressed in TH^+^ cells. For colocalization analyses, 6 coronal sections spanning 1.2 mm through the VTA from each of 4 rats were imaged using a Nikon laser scanning C2 system. Sections were surveyed at 4X magnification using a 594 nm excitation filter cube epifluorescence to identify the coronal section with the most expansive mCherry fluorescence. This coronal section was then imaged at 4X magnification with 405 nm, 488 nm, and 561 nm lasers to capture a 3 channel (DAPI, mCherry, TH) image of the entire section. Then, 2-4 images per hemisphere were taken at 40X magnification to again capture a 3-channel image (∼300×300 μm) for colocalization analyses.

#### Experiment 8: Effect of CNO on evoked excitatory postsynaptic currents in nucleus accumbens medium spiny neurons innervated by VTA dopamine terminals that express designer receptors (in vitro intracellular electrophysiological recordings)

Male, TH::Cre rats (n=10) received stereotaxic surgery to deliver VTA microinfusions of 1 μl of AAV8-hSyn-DIO-hM4D(Gi)-mCherry (n=5) or AAV8-hSyn-DIO-hM3D(Gq)-mCherry (n=5). After 4-6 weeks to allow the receptors to be expressed in striatal dopamine terminals, rats were anaesthetized with isoflurane and decapitated. Brains were rapidly extracted and submerged in an ice-cold HEPES-based artificial cerebrospinal fluid (ASCF) solution containing (in mM) 92 NaCl, 2.5 KCl, 1.2 NaH_2_PO_4_, 20 HEPES, 30 NaHCO_3_, 25 glucose, 5 sodium ascorbate, 2 thiourea, 3 sodium pyruvate, 12 N-acetyl-L-cysteine (NAC), 10 MgSO_4_, and 0.5 CaCl_2_ (pH adjusted to ≈7.3-7.4 using 10 M NaOH) saturated with 95% O_2_/5% CO_2_. Coronal slices (300 μm) containing the nucleus accumbens were obtained using a vibratome (Leica VT1200). Slices were transferred to a high-choline recovery solution containing (in mM): 92 choline chloride, 2.5 KCl, 1.2 NaH_2_PO_4_, 30 NaHCO_3_, 20 HEPES, 25 glucose, 5 sodium ascorbate, 2 thiourea, 3 sodium pyruvate, 12 NAC, 10 MgSO_4_, and 0.5 CaCl_2_, where they were allowed to recover for 12 min at 34°C. Subsequently, slices were transferred to normal ACSF solution containing (in mM): 124 NaCl, 5 KCl, 1.25 NaH_2_PO_4_, 2 MgSO_4_, 26 NaHCO_3_, 2 CaCl_2_, and 10 dextrose, where they recovered for at least one hour at 20-22°C. Once transferred into the recording chamber, slices were perfused with normal ACSF at 2 ml/min and were visualized using an upright fluorescence microscope with a 40X water-immersion objective, differential interference contrast optics (Olympus BX51WI), and XM-10 monochrome camera for viewing (Olympus, CellSens V1.8). Infection of the nucleus accumbens core was verified at 4X and 40X magnification prior to recordings (see Fig. S8).

Whole-cell patch clamp pipettes made using boroscillate glass (1.0 mm OD, 3 – 5 MΩ) were filled with a recording solution containing (in mM): 140 K-gluconate, 5 NaCl, 2 MgCl_2_, 10 HEPES, 0.5 EGTA, 2 ATP–tris, 0.4 GTP–tris (pH adjusted to 7.25 using KOH; 270 – 280 mOsm) and were lowered onto visually-identified neurons in the nucleus accumbens core, and tight seals were obtained (1.3 −6.6 GΩ). Cells were allowed to stabilize in whole cell configuration for 10 min prior to recordings. Access resistance was 19.9 ± 2.2 mΩ, and series resistance was uncompensated. Recordings were obtained using a Multiclamp 700B amplifier (Molecular Devices), digitized (Digidata 1440A, Molecular Devices), and were stored using pClamp 10.3 software (Molecular Devices). All cells recorded had a resting membrane potential below −65 mV. Cellular input resistance, membrane capacitance, access resistance were monitored during each recording condition.

Cells were first selected based on visual identification; medium spiny neurons and GABAergic interneurons possess smaller soma than cholinergic interneurons (8-20 μm vs 20-50 μm^36,83,84^) and any cells with soma >30 μm were not recorded from. GABAergic interneurons and medium spiny projections neurons of the accumbens core were differentiated by injecting 500 ms hyperpolarizing and depolarizing current steps between −100 and 100 pA in 10 pA intervals from the holding potential of −70mV. Peak input resistance was measured at the largest voltage change in response to −100 pA pulse, and steady state input resistance was assessed just prior to the end of the current step. Action potential properties were measured from the first action potential evoked in response to a positive current injection.

Synaptic responses were evoked using a bipolar stimulating electrode made from two tungsten electrodes (1 MΩ, FHC Inc.) placed approximately 30 µm from the recording electrode. Evoked AMPA-receptor-mediated excitatory postsynaptic currents (EPCSs) were recorded at a −70 mV, near the resting membrane potential of medium spiny neurons of the accumbens core^85^ using constant current stimulation pulses. For each cell, at least 10 consecutive synaptic responses free from artifacts or action potentials were averaged for each phase of the recordings. Dopamine release from VTA terminals to the nucleus accumbens core is under tonic inhibition from aspiny GABAergic interneurons^86^, and dopamine can also modify inhibition in the accumbens^85,87^. Because GABA receptors can influence the excitability of medium spiny neurons, we included picrotoxin (50 μM) in the ACSF to block GABA_A/C_-mediated inhibition and better assess the effects of CNO on VTA inputs to medium spiny neurons. Recordings were obtained before and after 5 min application of 1 μM CNO, and were also obtained after 20 min washout of CNO in the continued presence of picrotoxin. The amplitudes of averaged synaptic currents were measured using Clampfit 8.2 software (Molecular Devices) and normalized to the amplitude of responses recorded prior to CNO application.

## Histology and imaging

Rats from experiments 2-6 were deeply anesthetized with euthanyl^™^ (sodium pentabarbitol 240 mg/kg i. p.) and perfused with approximately 250 ml of 0.02 M phosphate buffered saline (PBS; pH 7.2), and 150 ml of 4% paraformaldehyde in 0.02 M PBS (pH 7.2). Immediately following perfusion, brains were removed and stored in 50 ml of a 4% paraformaldehyde 30% sucrose solution for cryoprotection for approximately 2-3 days. They were then removed from the sucrose solution and stored at −80°C until they were sectioned (40 μm thick) using a cryostat at −20°C. Nissl staining was conducted to assess histological placements of injector tips in the NAc core and shell in experiments 5 and 6. Unamplified mCherry signal was assessed to verify successful VTA designer receptor expression in experiments 2-6 by letting slides dry for 24 h after removal from the −20°C freezer and then coverslipping them with vectasheild^™^. For the purpose of verifying VTA expression of the designer receptor mCherry in experiments 2-6 we assessed mCherry fluoresecence using a Leica DM4000B epifluorescence microscope at 5X and 10X magnification.

## Analyses and Statistics

### Software

With the exception of electrophysiological data that were analyzed with SigmaPlot^™^ version 11, all statistical analyses were conducted using SPSS^™^ version 20. All analyses used an alpha level of *p*=.05. All graphs were made using SigmaPlot^™^ version 12. All significant interactions were pursued using Bonferroni-corrected t-tests. When sphericity was violated for the repeated measures analyses Huynh-Feldt corrections were applied.

### Home-cage alcohol exposure

The dependent measure for home-cage alcohol exposure was grams of ethanol consumed in 24 h as a function of rat weight in kilograms (g/kg). Ethanol consumption was analyzed using repeated measures (RM) ANOVA with Session (1, 2, 3…12) as the only factor, except for experiment 1 where Group (CS/NS, CS/noNS) was included as a factor.

### Pavlovian conditioning with context alternation and CS-triggered alcohol-seeking tests

For Pavlovian conditioning training and CS-triggered alcohol-seeking tests we measured the number of port entries made during each CS trial and during the 10 s intervals preceding each CS trial (PreCS). Total port entries are all port entries made during a session. For Pavlovian conditioning, PreCS and CS port entries in the alcohol context were compared to port entries made during the PreNS and NS in the neutral context. Port entries made across the acquisition of Pavlovian conditioning were analyzed in a Session (1, 2, 3, …12) by Context (Alcohol, Neutral) by Interval (PreCS/PreNS, CS/NS) RM ANOVA. The total number of port entries made in each training session was analyzed in a Session (1, 2, 3, …12) by Context (Alcohol, Neutral) RM ANOVA.

For CS-triggered alcohol-seeking tests, PreCS and CS port entries were compared to each other and across contexts. The RM ANOVA conducted for the CS-triggered alcohol-seeking tests included the factors Context (Alcohol, Neutral) and Interval (PreCS, CS). Port entries made outside of the CS interval (NonCS port entries; i.e., Total minus CS port entries) were analyzed using a RM ANOVA with the factor Context (Alcohol, Neutral).

For experiments assessing CS-triggered alcohol-seeking in the neutral context, test data were analyzed in a within-subjects, Interval (PreCS, CS) by Treatment (Saline, SCH23390, Eticlopride) (Vehicle, 10 mg/kg CNO, 20 mg/kg CNO) (Vehicle, 10mg/kg CNO, .1 mg/kg Clozapine) (Vehicle, 10 mg/kg CNO) RM ANOVA. NonCS port entries were also analyzed in a RM ANOVA with the within-subjects factor Treatment.

For microinfusion experiments, test data were analyzed in a Treatment (Vehicle, CNO) by Context (Alcohol, Neutral) by Interval (PreCS, CS) RM ANOVA. NonCS port entries were analyzed in a RM ANOVA with the factors Treatment (Vehicle, CNO) and Context (Alcohol, Neutral).

### Repeated test sessions and reinstatement in the alcohol and neutral contexts

For clarity, repeated test session and reinstatement test data are shown as normalized CS port entries (CS port entries minus PreCS port entries). Normalized CS port entries made across repeated test sessions were analyzed in a Session (1, 2, 3, 4) by Context (Alcohol, Neutral) RM ANOVA.

Normalized CS port entries made during the last repeated test session and reinstatement test were analyzed in a Reinstatement Context (Alcohol, Neutral) by Phase (Last test, Reinstatement Test) between-subjects repeated measures ANOVA.

### Electrophysiology analyses

RM ANOVAs were used to assess changes in synaptic response and cellular properties induced by drug application. The data met the requirements for normality (Kolmogorov-Smirnov test with Lilliefors’ correction).

## Supporting information

## Data Availability

The data from these experiments will be made available from the corresponding authors upon request.

## Acknowledgements

We acknowledge Steve Cabilio for support with Med-PC programming and data extraction and Karl Deisseroth and his laboratory for the donation of founder TH::Cre rats. This work was supported by grants from CIHR (MOP-137030, N.C.), FRQS (Chercheur Boursier Junior 2), CFI (John R. Evans Leaders Fund), NSERC (387 224-2010, N.C.; 2014-0507, A.C.).

## Author contributions

N.C. and M.V designed the behavioral experiments. N.C., M.V., and I.T. designed the microscopy co-localization experiment. A.C. and I.G. designed, conducted, and analyzed the in vitro electrophysiology experiments. M.V. analyzed all the behavioral and co-localization data. M.V., A.Z., and S.L., conducted all behavioral experiments and breeding of TH::Cre rats. I.T. assisted with breeding TH::Cre rats. M.V., I.G., A.C., and N.C. wrote the manuscript. All authors reviewed the manuscript and provided comments.

