## Supplementary material for "Divergent mesolimbic dopamine circuits support alcohol-seeking triggered by discrete cues and contexts"

| Experiment | n | Figure(s) | Description |
|---|---|---|---|
| 1 | 22 | 1 | An alcohol context has an enduring impact on alcohol-seeking triggered by a discrete cue and amplifies relapse. |
| 1b | 11 | 2b | Port-entries made between CS trials are attenuated by systemic administration of dopamine D1 and D2 receptor antagonists, and CS-triggered alcohol-seeking is attenuated only by a D2 receptor antagonist. |
| 2 | 12 | 2c, S5a | CS-triggered alcohol-seeking is attenuated by chemogenetically silencing VTA dopamine neurons. |
| 3 | 13 | 2d, S5b | CS-triggered alcohol-seeking is unaffected by clozapine-n-oxide and clozapine. |
| 4 | 18 | S6 | CS-triggered sucrose-seeking is unaffected by chemogenetically silencing VTA dopamine neurons. |
| 5 | 8 | 3b, 3d, 3e, S5c | Silencing a dopaminergic VTA-to-NAc core circuit attenuated CS-triggered alcohol-seeking in an alcohol context and a neutral context. |
| 6 | 11 | 3c, 3f, 3g, S5d | Silencing dopaminergic VTA-to-NAc shell projection attenuated CS-triggered alcohol seeking in an alcohol context, but not in a neutral context. |
| 7 | 4 | 4a-d | Colocalization analysis of specificity and transduction efficiency. |
| 8 | 10 | 4e-j | Chemogenetic control of dopaminergic activity modulates striatal medium spiny neuron activity. |

**Supplementary Table 1│** For use when referring to online methods and supplementary figure captions, experiments are listed by number, and the sample size (n), relevant Figure numbers, and brief descriptions are provided.

| Group | Alcohol Context | Neutral Context | CS | NS | # of Rats by experiment | | | | | |
|---|---|---|---|---|---|---|---|---|---|---|
| | | | | | 1 | 2 | 3 | 4 | 5 | 6 |
| CS/NS | 1 | 2 | White Noise | Clicker | 3 | 4 | 1 | 5 | 0 | 4 |
| | 1 | 2 | Clicker | White Noise | 2 | 1 | 4 | 4 | 3 | 1 |
| | 2 | 1 | White Noise | Clicker | 3 | 4 | 5 | 4 | 3 | 4 |
| | 2 | 1 | Clicker | White Noise | 3 | 3 | 3 | 5 | 2 | 2 |
| CS/NoNS | 1 | 2 | White Noise | NA | 2 | | | | | |
| | 1 | 2 | Clicker | NA | 3 | | | | | |
| | 2 | 1 | White Noise | NA | 3 | | | | | |
| | 2 | 1 | Clicker | NA | 3 | | | | | |
| Total n | | | | | 22 | 12 | 13 | 18 | 8 | 11 |

**Supplementary Table 2**│Breakdown of the number of rats from each experiment that received the different combinations of discrete, auditory stimuli used as the conditioned stimulus (CS) or neutral stimulus (NS) in each of the 2 context configurations used as either the alcohol context or neutral context. Context 1 had dark walls, a smooth polycarbonate floor, and a lemon odour. Context 2 had clear walls, a wire-mesh floor, and an almond odour.

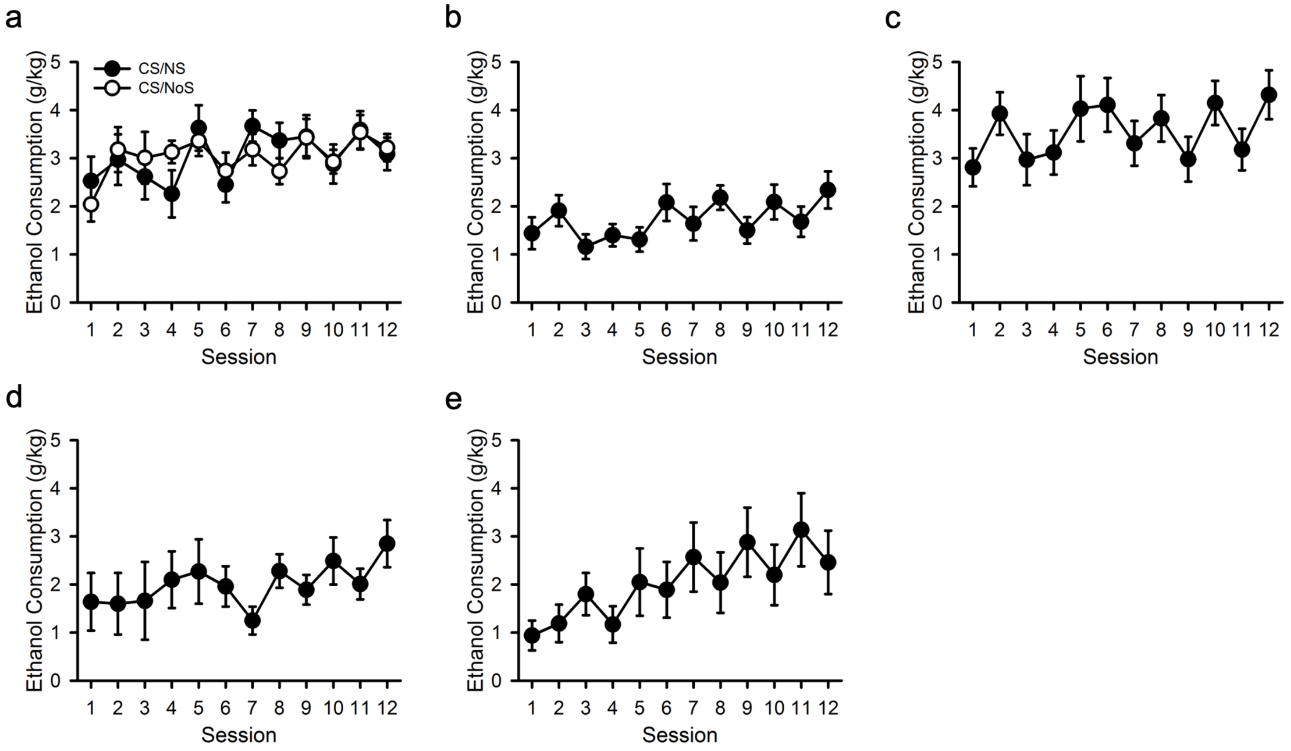


**Supplementary Fig. 1│ Alcohol consumption and preference increase over intermittent access home-cage ethanol exposure.** Rats received access to a 15% ethanol solution in their home-cage every other day, with water continuously available.

**a**, Ethanol consumption in wild-type rats increased across sessions, but did not differ between groups in Experiment 1 [Session, $F_{(11, 220)}$ = 3.875, $p$<.001; Group, $F_{(1, 20)}$ <.001, $p$=.993; Session x Group $F_{(11, 220)}$ = 1.106, $p$=.358].

Ethanol consumption increased across sessions in TH::Cre$^{+/-}$ rats in:

**b**, Experiment 2, [$F_{(11, 132)}$=3.799, $p$<.001].

**c**, Experiment 3, [$F_{(11, 154)}$=4.139, p<.001].

**d,** Experiment 5, [$F_{(11, 77)}$=2.257, $p$=.019].

**e**, Experiment 6, [$F_{(11, 110)}$=7.140, $p$<.001].

All averaged data are shown as mean ± s.e.m.

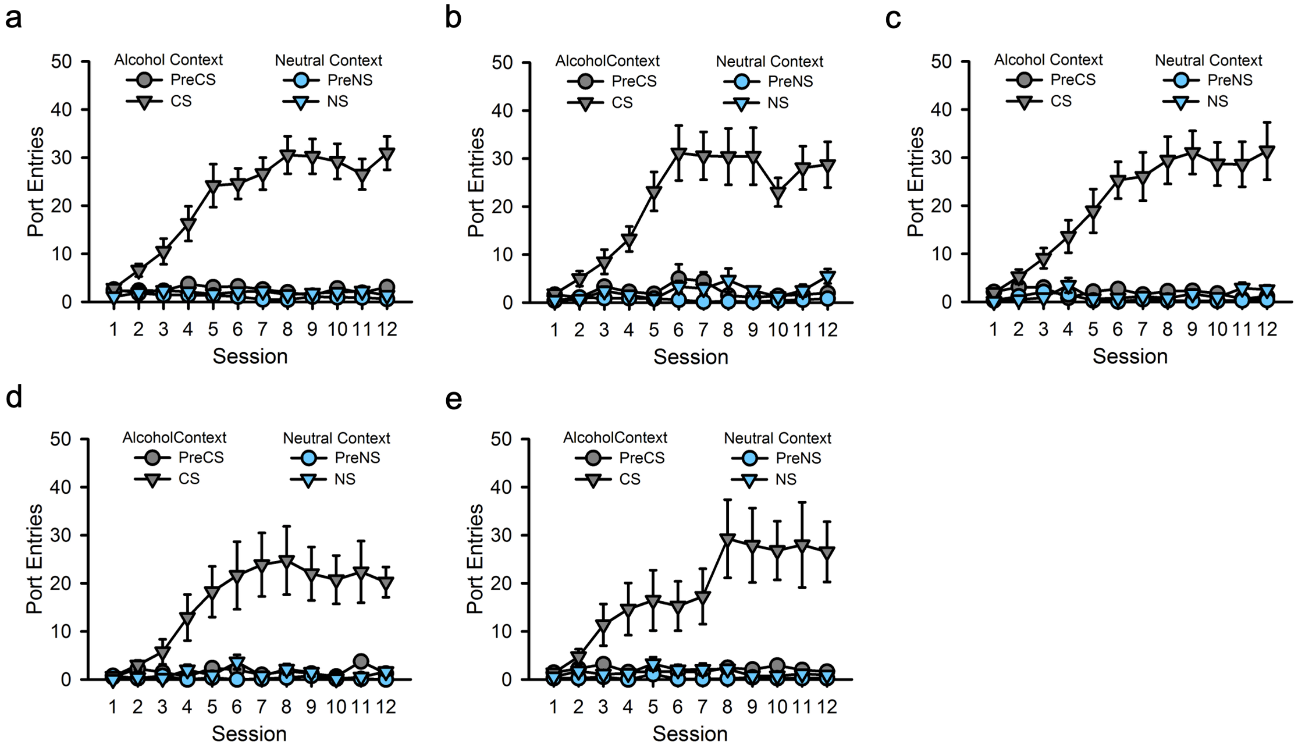


**Supplementary Fig. 2│Acquisition of Pavlovian conditioning with context alternation.** Rats received Pavlovian conditioning sessions every other day in a distinct ‘alcohol context’ where a discrete auditory conditioned stimulus (CS) was paired with 15% ethanol. On alternating days, rats were exposed to a different, ‘neutral context’ where a distinct, neutral auditory stimulus (NS) was presented without alcohol.

**a**, In experiment 1, while in the neutral context, half of the rats received presentations of an auditory stimulus that was distinct from their CS (referred to as the neutral stimulus, NS group), whereas the remainder did not (noNS group; **Tables S1 and 2**). The purpose for having 2 groups was to test whether prior results obtained using this paradigm could be attributed to differences in acoustical salience across the two contexts due to the the lack of an auditory stimulus in the neutral context. Since there were no significant main effects or interactions as a function of this condition, the data reported here were collapsed across groups. In all other experiments an NS was played in the neutral context during training. CS-triggered fluid port entries increased into a plateau over Pavlovian conditioning sessions, whereas port entries during the NS or during an equivalent period of time immediately before either stimulus (PreCS and PreNS) remained low. Experiment 1 [Session x Context x Interval, $F_{(11, 220)}$=17.37, $p$<.001].

**b**, Experiment 2 [Session x Context x Interval $F_{(11, 121)}$=8.876, $p$<.001].

**c**, Experiment 3 [Session x Context x Interval, $F_{(11, 132)}$=11.323, $p$<.001].

**d**, Experiment 5 [Session x Context x Interval, $F_{(11, 77)}$=5.352, $p$<.001].

**e**, Experiment 6 [Session x Context x Interval, $F_{(11, 110)}$=8.149, $p$<.001].

All averaged data are shown as mean ± s.e.m.

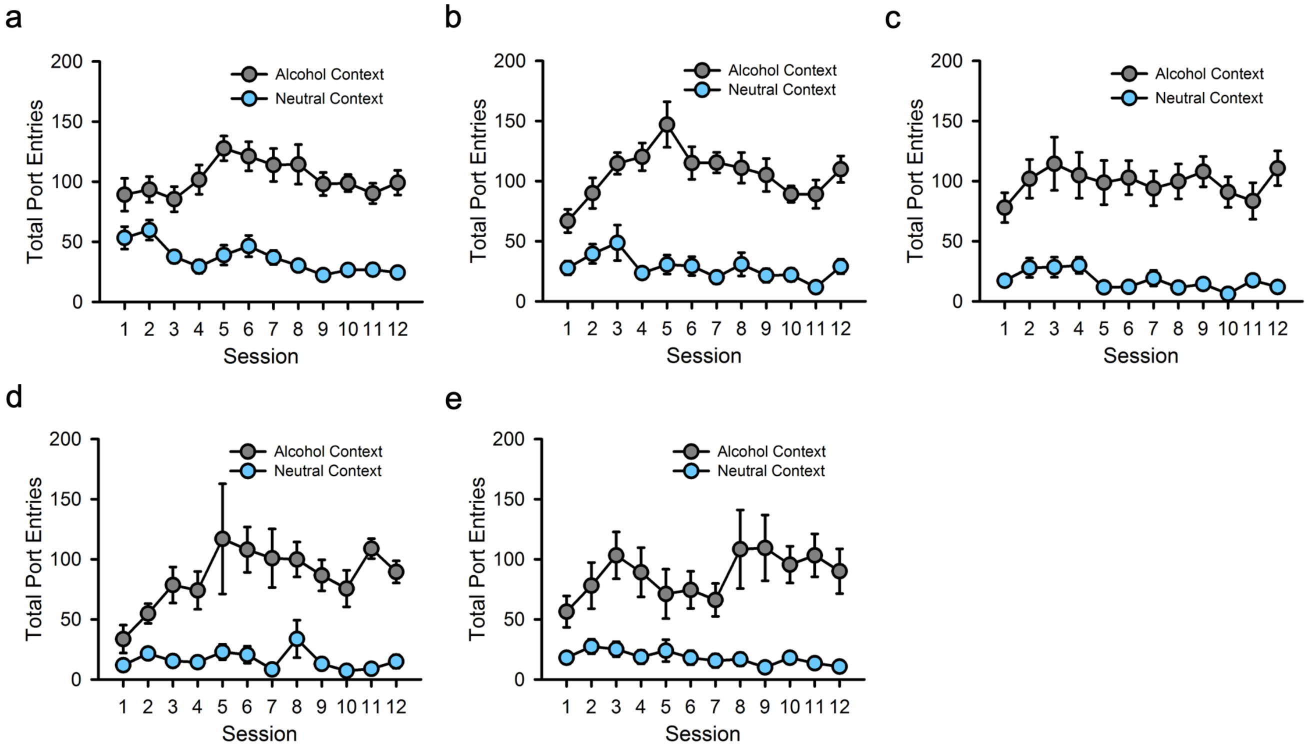


**Supplementary Fig. 3│Total port entries during acquisition of Pavlovian conditioning with context alternation.** Rats received Pavlovian conditioning sessions every other day in a distinct ‘alcohol context’ where a discrete auditory conditioned stimulus (CS) was paired with 15% ethanol. On alternating days, rats were exposed to a different, ‘neutral context’ where a distinct, neutral auditory stimulus (NS) was presented without alcohol. Total port entries were elevated in the alcohol context and typically waned across session in the neutral context.

**a,** Experiment 1 [Context, $F_{(1, 20)}$=100.59, $p$<.001; Context x Session, $F_{(11, 220)}$=3.45, $p$<.001].
**b**, Experiment 2 [Context, $F_{(1, 11)}$=129.409, $p$<.001; Context x Session, $F_{(11, 121)}$=3.054, $p$=.001].
**c**, Experiment 3 [Context, $F_{(1, 12)}$=42.715, $p$<.001; Context x Session, $F_{(11, 132)}$=13.480, $p$<.001].
**d**, Experiment 5 [Context, $F_{(1, 7)}$=47.187, $p$<.001; Context x Session, $F_{(11, 77)}$=1.605, $p$=.114].
**e**, Experiment 6 [Context, $F_{(1, 10)}$=25.726, $p$<.001; Context x Session, $F_{(11, 110)}$=2.235, $p$=.017].
All averaged data are shown as mean ± s.e.m.

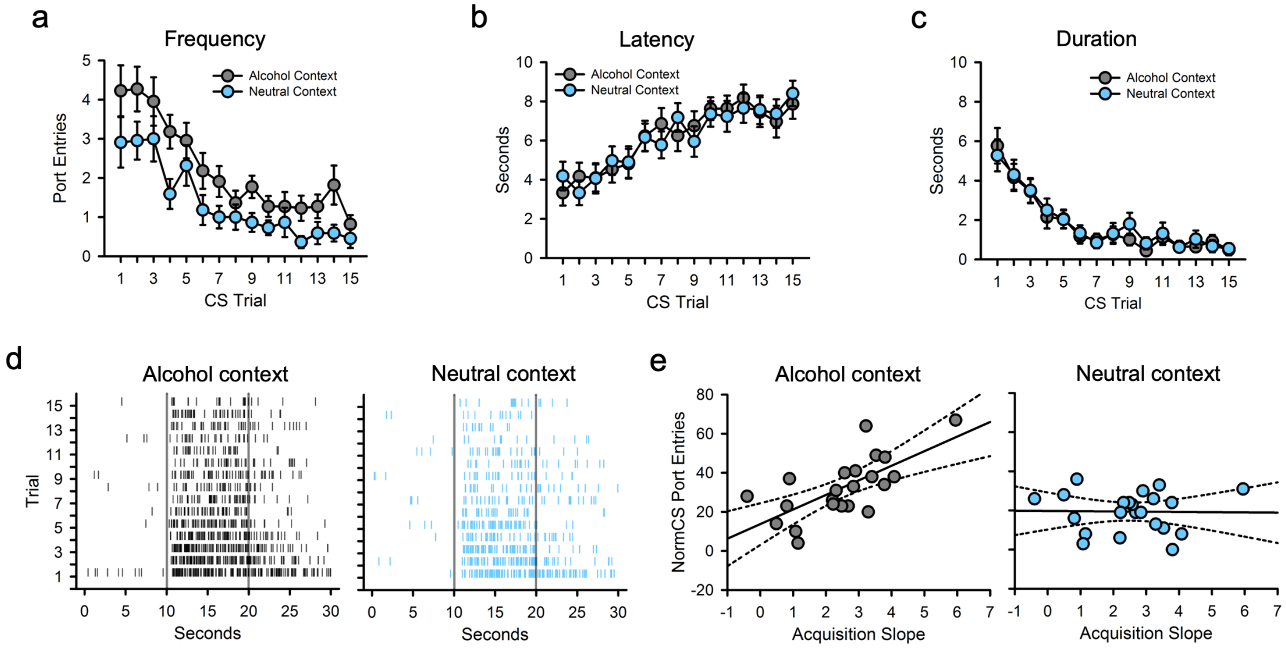


**Supplementary Fig. 4│Analysis of the conditioned response form and predictors of CS-triggered alcohol-seeking.** We analyzed the frequency, latency, and duration of CS-triggered port entries at test on a trial-by-trial basis, as well as the predictors of CS-triggered alcohol-seeing.

**a**, CS-triggered port entries decreased across CS trials at test, but were elevated overall in the alcohol context [Context, $F_{(1, 20)}$ = 13.53, $p$<.001; Trial, $F_{(14, 280)}$=16.243, $p$=.001; Trial x Context, $F_{(14, 280)}$=.530, $p$=.915].

**b**, The latency to make a CS-triggered port entry increased comparably across CS trials at test in both contexts [Trial, $F_{(14, 280)}$=12.329, $p$<.001; Context, $F_{(1, 20)}$=.006, p=.939; Context x Trial, $F_{(14, 280)}$= .488, $p$=.939].

**c**, The duration of CS-triggered port entries decreased comparably across CS trials at test in both contexts [Trial, $F_{(14, 280)}$=19.79, $p$<.001; Context, $F_{(1, 20)}$=.169, $p$=.685; Context x Trial, $F_{(14, 280)}$= .279, $p$=.996].

**d**, Raster plots showing every port entry made during the PreCS (0-10 s), CS (10-20 s) and PostCS (20-30 s) intervals by all rats at test in both contexts.

**e**, We used a stepwise linear regression procedure to identify the model that best predicted CS-triggered port entries at test in the alcohol context (**Table S3**) and neutral context (**Table S4**). The following variables were considered: alcohol intake (g/kg) on sessions 1 and 12 of home-cage alcohol exposure, the slope of alcohol intake over all 12 home-cage alcohol exposure sessions, normalized CS port entries on sessions 1 and 12 of Pavlovian conditioning, and the slope of acquisition over all 12 Pavlovian conditioning sessions. Rate of acquisition of CS port entries during training predicted normalized CS-triggered port entries (i.e., CS minus PreCS port entries) at test in the alcohol context [$F_{(1, 20)}$=17.587, $p$<.001; $r^2$=.468], but not the neutral context ($p$ =*n.s.*). Dotted lines are 95% confidence intervals. Solid lines are regression lines. All averaged data are shown as mean ± s.e.m.

| Retained Variable | β | Std Error | *t* | *p* |
|---|---|---|---|---|
| Pavlovian Acquisition Slope | 7.467 | 1.780 | 4.194 | <.001 |
| Excluded Variables | | | | |
| Alcohol Intake Slope | .057 | | .340 | .737 |
| NormCS Port Entries Session 1 | .113 | | .654 | .521 |
| NormCS Port Entries Session 12 | .165 | | .659 | .518 |
| Alcohol Intake Session 1 | .101 | | .602 | .554 |
| Alcohol Intake Session 12 | .270 | | 1.717 | .102 |

**Supplementary Table 3**│Breakdown of the parameters retained and excluded from a stepwise linear regression predicting Normalized CS-triggered port entries made at test in the alcohol context (related to Fig. S4e).

| Excluded Variables | β | Std Error | *t* | *p* |
|---|---|---|---|---|
| Pavlovian Acquisition Slope | -2.211 | | -.770 | .454 |
| Alcohol Intake Slope | -35.941 | | -1.302 | .213 |
| NormCS Port Entries Session 1 | -.513 | | -.881 | .392 |
| NormCS Port Entries Session 12 | .206 | | .743 | .469 |
| Alcohol Intake Session 1 | .313 | | .154 | .880 |
| Alcohol Intake Session 12 | 3.603 | | 1.254 | .229 |

**Supplementary Table 4**│Breakdown of the parameters excluded from a stepwise linear regression predicting Normalized CS-triggered port entries made at test in the neutral context (related to Fig. S4e).

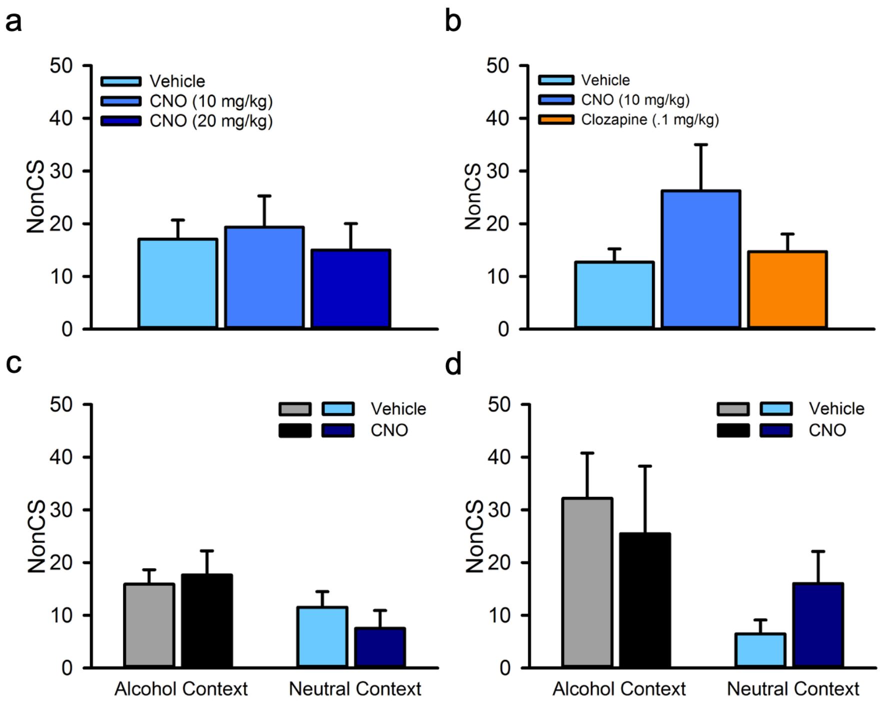


**Supplementary Fig. 5│NonCS port entries made during test for CS-triggered alcohol-seeking.** Port entries made between CS trials (NonCS) are a measure of general port directed behavior.

**a**, Experiment 2: NonCS port entries made in the neutral context by TH::Cre rats transfected with hSyn-DIO-hM4Di-mCherry were similar across tests [Treatment, $F_{(2, 22)}$=.288, $p$=.752].

**b**, Experiment 3: NonCS port entries made in the neutral context by TH::Cre rats transfected with hSyn-DIO-mCherry were similar across tests [Treatment, $F_{(2, 24)}$=1.678, $p$=.208].

**c**, Experiment 5: NonCS test port entries made by TH::Cre rats that were transfected with hSyn-DIO-hM4Di-mCherry and received microinfusions of CNO or vehicle in the NAc core were similar across context and treatment conditions [Context, $F_{(1, 7)}$=4.307, $p$=.077; Treatment, $F_{(1, 7)}$=.063, $p$=.809; Context by Treatment, $F_{(1, 7)}$=.688, $p$=.434].

**d**, Experiment 6: NonCS port entries made by TH::Cre rats that were transfected with hSyn-DIO-hM4Di-mCherry and received microinfusions of CNO or vehicle in the NAc shell were similar across treatments, but elevated in the alcohol context relative to the neutral context [Treatment, $F_{(1, 10)}$=.03, $p$=.866; Context, $F_{(1, 10)}$=16.221, $p$=.002].

All averaged data are shown as mean ± s.e.m.

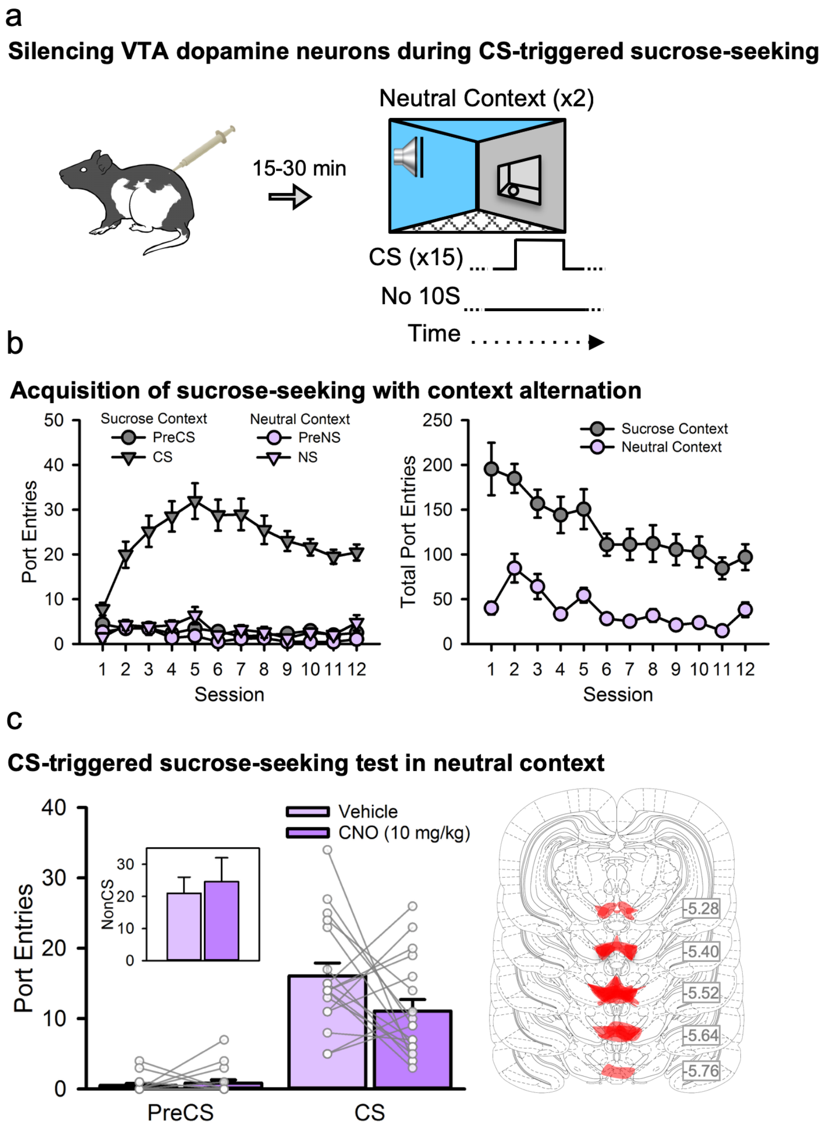


**Supplementary Fig. 6│Silencing VTA dopamine neurons failed to attenuate CS-triggered sucrose-seeking.** We examined the necessity of VTA dopamine neurons for CS-triggered sucrose-seeking.

**a**, Using naïve TH::Cre rats (n=18) we conducted Pavlovian conditioning with context alternation as described in **Fig. 1a**, except that CS trials were paired with 10% sucrose. All rats were tested for CS-triggered sucrose-seeking in a neutral context after chemogenetic silencing of VTA dopamine neurons.

**b**, CS port entries increased into a plateau across conditioning sessions whereas port entries during the NS or preceding either interval (PreCS, PreNS) remained stably low [Session x Context x Interval, $F_{(11, 187)}$=5.140, $p$<.001]. Total port entries waned across sessions but decreased to lower levels in the neutral context [Context, $F_{(1, 17)}$=64.549, $p$<.001; Context x Session, $F_{(11, 187)}$=3.215, $p$<.001].

**c**, CS-triggered port entries were elevated relative to PreCS port entries and were similar for vehicle and 10 mg/kg CNO [Interval by Context, $F_{(1, 17)}$=3.691, $p$=.072]. The inset graph shows NonCS port entries, which were similar across both tests [Treatment, $F_{(1, 17)}$=.173, $p$=.682]. Histology sections (right) depict maximal mCherry expression in the midbrain for every rat. All averaged data are shown as mean ± s.e.m.

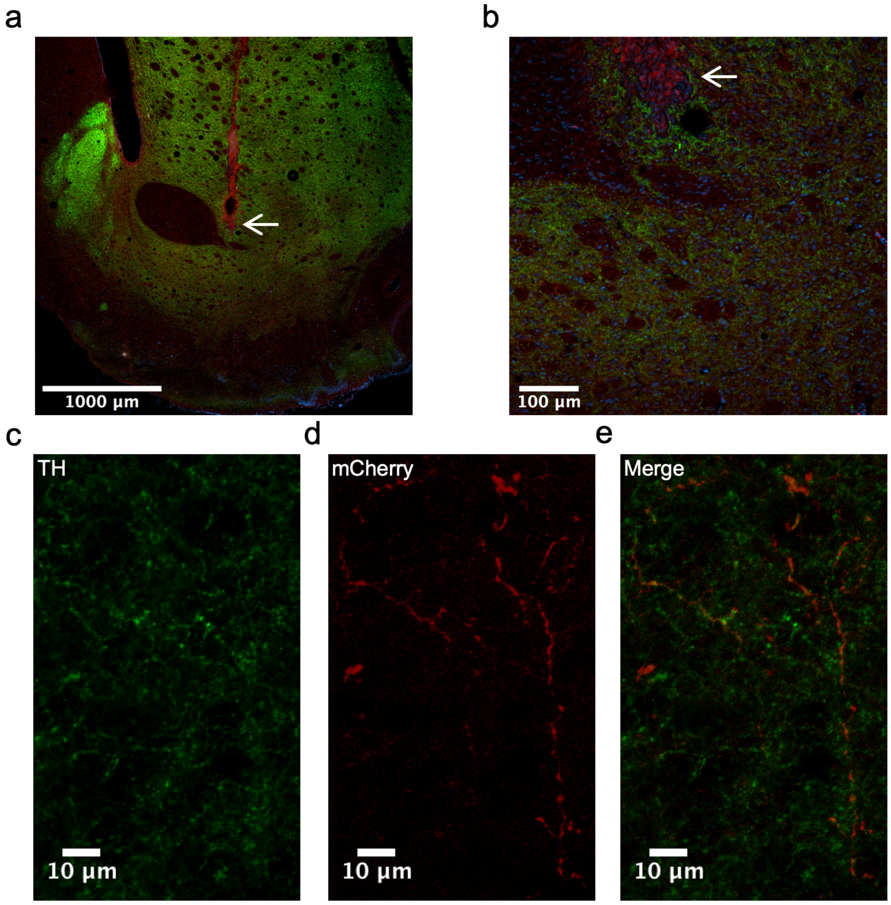


**Supplementary Fig. 7│Immunofluorescence images from the nucleus accumbens (NAc) core.**

**a**, A 4X image of the ventral striatum showing injector placement (arrow) in the NAc core.

**b**, A 40X image showing the lower-right quadrant of NAc nucleus accumbens core.

**c**, A 60X image showing TH+ neural processes in the NAc core near the injector tip in the previous panels.

**d**, The same processes showing mCherry expression (reporter for DIO-hM4Di designer receptor).

**e**, A merge of TH and mCherry expression showing overlap.

All images in this figure are z-projections of maximum signal to highlight processes across the z-plane.

DAPI is blue, TH is green, and mCherry is red.

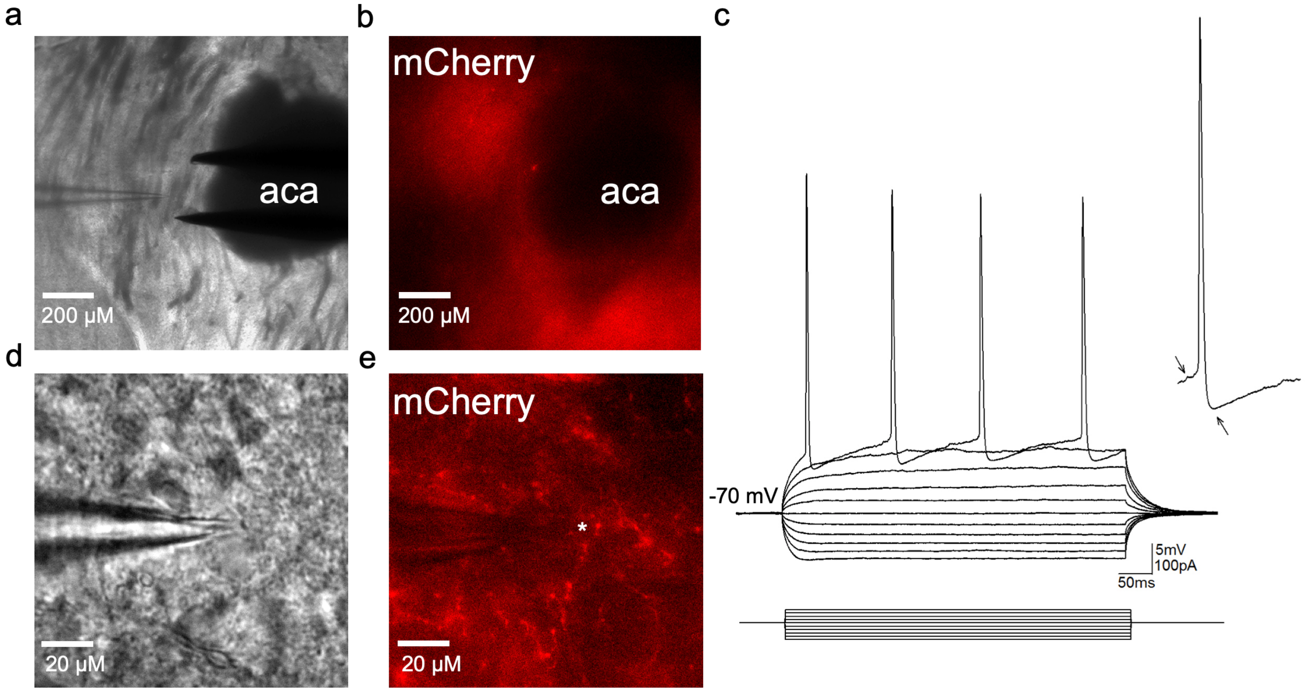


**Supplementary Fig. 8│Microscopy images obtained during electrophysiological recordings from nucleus accumbens (NAc) medium spiny neurons (MSN).**

**a**, A representative brightfield image showing the position of stimulating and recording electrodes in the NAc core.

**b**, A representative fluorescence image of the same coronal slice showing strong mCherry expression in VTA fibers spread throughout the NAc core (aca: anterior portion of the anterior commissure).

**c**, Superimposed membrane potential responses in a representative MSN in response to hyperpolarizing and depolarizing current steps showing isolated action potentials. Arrows indicate the slow ramp-like depolarization prior to firing, and large fast after-hyperpolarization that are typical of MSN neurons. Resting membrane potential of MSNs was -66.1 ± .4 mV, capacitance was 23.4 ± 3.2 pF, and cellular input resistance was 188 ± 14.5 MΩ. A 5-min bath application of 1 μM CNO to inhibit hM4Di-expressing terminals of VTA neurons within the NAc core did not produce significant changes in action potential number (2.4 ± .5 at baseline vs. 1.7 ± .5 in CNO, $t$=1.18, $p$=0.28), width (9.7 ± .7 ms at baseline vs. 9.5 ± .7 ms in CNO, $t$=1.76, $p$=0.14), or height (93.9 ± 3.4 mV at baseline vs. 94.6 ± 3.9 mV in CNO, $t$=0.23, $p$=0.83) during injection of positive current steps.

**d**, Image of an MSN that was recorded from.

**e**, Image of the same MSN in **d** relative to mCherry-expressing dopaminergic VTA fibers.
